# Suppressing peatland methane production by electron snorkeling through pyrogenic carbon

**DOI:** 10.1101/2020.02.15.950451

**Authors:** Tianran Sun, Juan J. L. Guzman, James D. Seward, Akio Enders, Joseph B. Yavitt, Johannes Lehmann, Largus T. Angenent

## Abstract

Northern peatlands are experiencing more frequent fire events as a result of changing climate conditions. Forest fires naturally result in a direct and negative climate impact by emitting large amounts of carbon into the atmosphere. Recent studies show that this extensive emission may shift the soil carbon regime from a sink to a source. However, the fires also convert parts of the burnt biomass into pyrogenic carbon. Here, we show an indirect, but positive, climate impact induced by fire-derived pyrogenic carbon. We found that the accumulation of pyrogenic carbon reduced post-fire methane production from peatland soils by 13-24%. The conductive, capacitive, and redox-cycling electron transfer mechanisms enabled pyrogenic carbon to function as an electron snorkel, which redirected soil electron fluxes to facilitate alternative microbial respiration and reduced the rate of methane production by 50%. Given the fact that methane has a 34-fold greater warming potential than carbon dioxide, we estimate that global greenhouse gas emissions are reduced by 35 Tg CO_2_e annually through the electron snorkeling of pyrogenic carbon in peatlands. Our results highlight an important, but overlooked, function of pyrogenic carbon in neutralizing forest fire emissions and call for its consideration in the global carbon budget estimation.

## Introduction

Multiple lines of evidence show that forest fires become increasingly frequent in northern peatlands (i.e., latitude 40°–70°N) due to their vulnerability to a warming and drying climate^1,2^. Forest fires release large amounts of carbon into the atmosphere, but they also convert parts of the burnt biomass into pyrogenic carbon through the incomplete combustion of biomass. Due to its carbon enrichment and environmental persistence, pyrogenic carbon has shown the ability to buffer carbon emission during the fire^3^ and store carbon in peat soils from centuries to millennia after the fire^4^. In this context, current studies are striving to quantify the production of pyrogenic carbon caused by fires^5^ and fill up the storage gap in the global carbon budget^6^. However, thus far, it is still unclear how this long-term stored pyrogenic carbon interacts with the indigenous methane (CH_4_) production in peat soils.

Northern peatlands have historically been a sink for atmospheric carbon dioxide (CO_2_) but a source of atmospheric CH_4_^7,8^. Even though methanogens in peat soils are phylogenetically diverse, they produce CH_4_ mainly through hydrogenotrophic (Δ*G*^*0*^ = −33 kJ mol^-1^ hydrogen) and aceticlastic (Δ*G*^*0*^ = −36 kJ mol^-1^ acetate) methanogenesis^9^. The prevalence of either methanogenesis pathways depends on the nutrient level and feeding type (i.e., whether it is rain fed or stream fed) of a specific peat soil, but they all compete with anaerobic respiration that utilize alternative (other than oxygen) terminal electron acceptors^10^. Alternative terminal electron acceptors that have been reported in peat soils include manganese oxide, nitrate, iron minerals, sulfate, and dissolved and particulate organic matter in the order of high to low reduction potential (averagely from 0.8 to −0.2 V)^10,11^. Anaerobic respiration by *Geobacter* and *Shewanella* species with alternative terminal electron acceptors (hereafter referred to as alternative respiration) have been widely studied and demonstrated its favorability (Δ*G*^*0*^ = −40 to −234 kJ mol^-1^ hydrogen and −69 to −841 kJ mol^-1^ acetate) in substrate competition with methanogenesis, and thus suppressing CH_4_ production^12-14^. These microbes can also use pyrogenic carbon as an intermediate electron acceptor and conduit.

Pyrogenic carbon consists of electrically conductive carbon matrices and redox-active quinone and phenol functional groups^15,16^. The carbon matrices possess a wide 1.5 V potential range to transfer electrons to a variety of terminal electron acceptors^16^. Electron accepting and donating cycles of the quinone and phenol functional groups have been shown to mediate biogeochemical reactions in catalyzing microbial mineral reduction^17^ and enhancing organic contaminant transformation^18^. These demonstrated abilities of taking and passing electrons from and to different environmental substances led us to hypothesize that: (1) the accumulation of pyrogenic carbon can redirect soil electron fluxes and facilitate alternative respiration; and (2) the enhanced alternative respiration outcompetes methanogenesis and subsequently reduces CH_4_ production in peat soil. We propose to refer to the redirection of electron fluxes by the different electron transfer mechanisms of pyrogenic carbon as electron snorkeling^19^. To verify these hypotheses, we conducted 6 large-group bioelectrochemical and microcosm peat soil as well as pure-culture incubations, which in total contained 144 individual incubations to probe the functions of pyrogenic carbon in electron snorkeling and CH_4_ suppression. This systematic study helped to better understand the post-fire interactions as well as the underlying mechanisms between pyrogenic carbon and CH_4_ production, which will improve the estimates, and more importantly, the prediction of peatland gas emissions during a future of climate change.

## Results and Discussion

### Electron snorkeling suppressed methane production in peat soil

We tested the electron snorkeling of pyrogenic carbon and its effect on methanogenesis in bioelectrochemical and microcosm peat-soil incubations (**Supplementary Information (SI) Method S1-S4**). The soil was sampled from an ombrotrophic (i.e., rain fed) peat site that is known to produce CH_4_, with a field production rate of 3.5 Mg ha^-1^ yr^-1^ ref^20^, projecting to 119 Mg CO_2_e ha^-1^ yr^-1^. The CH_4_ production was very slow (<1 µmol g^-1^ soil carbon day^-1^) in the first few days of incubation. It started to rise after day 6. The CH_4_ production rate, thereafter, was in the range of 4.7-5.9 µmol g^-1^ soil carbon day^-1^ (or 0.08-0.1 mmol L^-1^ soil suspension day^-1^, see the pyrogenic carbon-free control treatment in **Figure 1** and **SI Figure S1**), which was in agreement with previously reported rates (0.02-0.1 mmol L^-1^ soil suspension day^-1^) from the same peat soil^21,22^. Hydrogenotrophic methanogen (*Methanoregula* and *Methanocellales*, with an average abundance of 1.2% for each) was found to dominate the peat-soil CH_4_ production (**SI Figure S2**).

**Figure 1.**
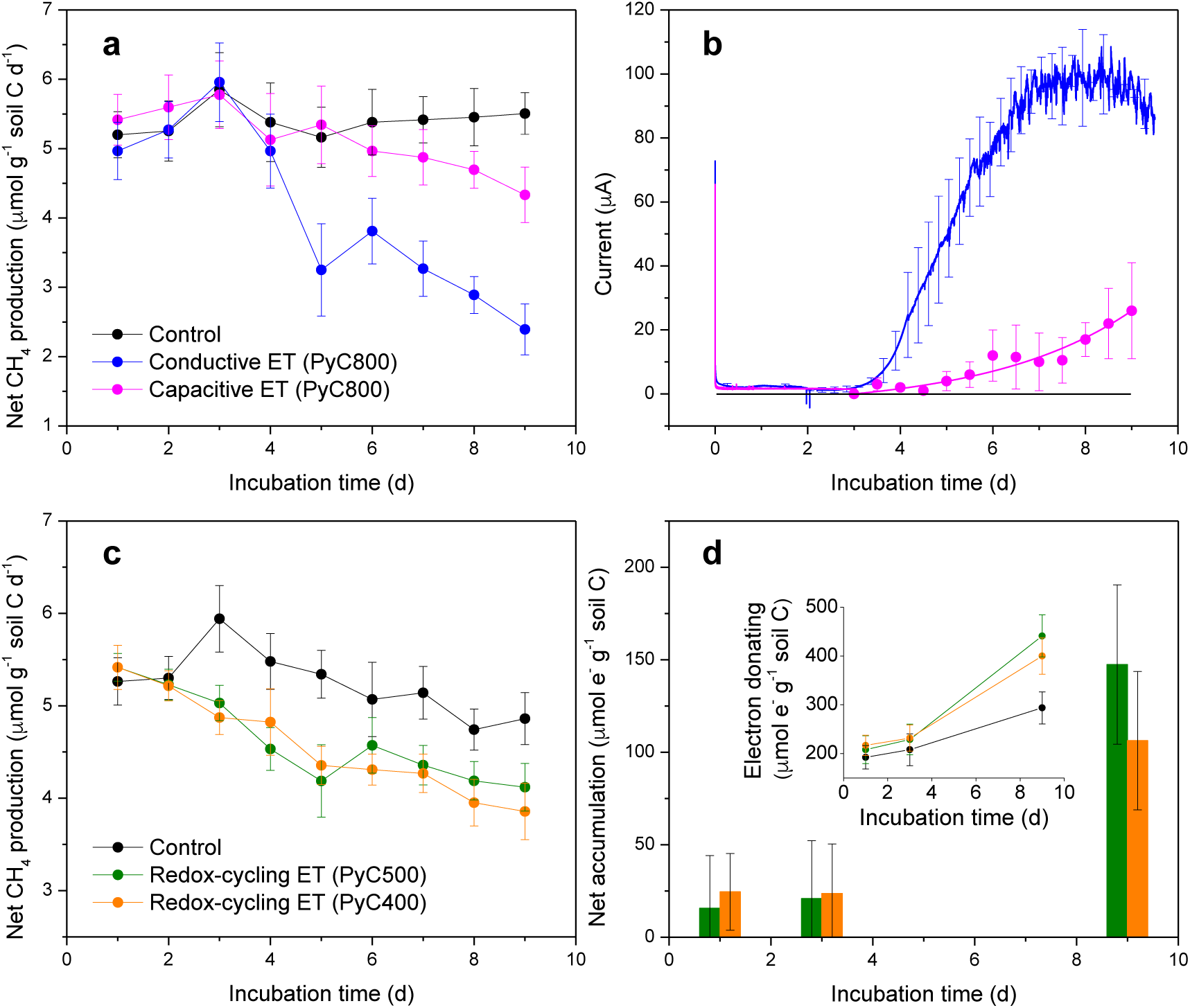
Electron snorkeling influenced CH_4_ production in the peat-soil incubations. **a.** Daily CH_4_ production rate in the bioelectrochemical peat-soil incubations that were associated with the conductive and capacitive electron transfer (ET) through the pyrogenic carbon matrices. Pyrogenic carbon that was produced at 800°C (PyC800) was used in the bioelectrochemical peat-soil incubations, with the application rate of 10 mg pyrogenic carbon g^-1^ soil. Figure legends in the chart **a** applies to the chart **b. b.** Current profiles generated from the conductive and capacitive ET through the pyrogenic carbon matrices. The intermittent current signals in the capacitive ET showed the highest current point of each discharging period at a 0.1 s^-1^ recording frequency. Full discharging current profile could be found at the chronoamperograms in **SI Figure S4. c.** Daily CH_4_ production rate in the microcosm peat-soil incubations that were associated with the redox-cycling ET of the pyrogenic carbon functional groups. Pyrogenic carbon that was produced at 400 and 500°C (PyC400 and PyC500) was used in the microcosm peat-soil incubations, with the application rate of 3 mg pyrogenic carbon g^-1^ soil. Figure legends in the chart **c** applies to the chart **d. d.** Net electron accumulation caused by the redox-cycling ET in the peat soil at the incubation day 1, 3, and 9. The net electron accumulation was calculated based on the electron donating difference (inset chart) between the pyrogenic carbon-amended peat soil and the control peat soil. Day 0 in the x axes in **a**-**d** indicate the time of adding pyrogenic carbon into the peat soil.

Pyrolyzed biomass was added to the peat soil (after 6 days of pre-incubation) to recreate the natural accumulation of pyrogenic carbon after vegetation fires. To distinguish pyrogenic carbon from native soil carbon, the original biomass was isotopically labelled with ^13^C, which resulted in a δ^13^C of pyrogenic carbon at 774±2.3‰. Even though the addition rate varied among incubations (0, 0.03, 3, and 10 mg pyrogenic carbon g^-1^ soil), we normalized the number of snorkeled electrons to a unit mass of either pyrogenic carbon or soil carbon to focus on the electron snorkeling efficiency instead of the effect of different addition rates. We found that three electron transfer mechanisms contributed to electron snorkeling and caused significant decreases in CH_4_ production: (1) conductive electron transfer through the carbon matrices; (2) capacitive electron transfer through the carbon matrices; and (3) redox-cycling electron transfer by the quinone and phenol functional groups (**Figure 1**).

The highest amount of CH_4_ suppression occurred in the bioelectrochemical peat-soil incubation that was associated with conductive electron transfer. Its total CH_4_ production (37±4.1 µmol g^-1^ soil carbon, blue line in **Figure 1a**) was 24% less than the pyrogenic carbon-free control treatment (49±3.8 µmol g^-1^ soil carbon, black line in **Figure 1a**). Pyrogenic carbon is able to conductively transfer electrons due to the inherent electrical conductivity of its carbon matrices^16,23^, which is derived from the overlapping of π-electron orbitals (i.e., π-electron system) among the polyaromatic carbon ring structures^24,25^. Conductive electron transfer often results in a continuous flow of electric current across environmental interfaces^26^. Following this principle, we implemented a bioelectrochemical circuit (**SI Figure S3a**) in the peat soil and successfully captured the current signals (red line in **Figure 1b**) while the conductive electron transfer was occurring. By integrating the current with the incubation time, we determined that the conductive electron transfer continuously snorkeled 785±120 µmol e^-^ g^-1^ soil carbon to alternative terminal acceptor (most likely the soil organic matter, **SI Method S1** and ref^9^) for facilitating alternative respiration. *Geobacter* spp. (with an average abundance of 1%) was determined as the major microbial member that performed alternative respiration (**SI Figure S2**). This number of snorkeled electrons was close to the reported electron accepting capacities (a few hundred to thousand µmol e^-^ g^-1^ soil carbon) of soil organic matter in several northern peat soils^27-29^, indicating the effectiveness of the conductive electron transfer in redirecting electron fluxes in peat soil.

In addition to continuous electron snorkeling that was induced by the conductive electron transfer, we found that the carbon matrices were also able to snorkel electrons intermittently through a series of capacitive electron storage and release cycles (i.e., capacitive electron transfer) as shown in the bioelectrochemical peat-soil incubation (red line in **Figure 1b**). During the capacitive electron transfer, electrons that were donated from alternative respiration were first stored in the π-electron system of the carbon matrices^30^. These stored electrons were surrounded and charge balanced by protons (the co-products with electrons when *Geobacter* dominates the alternative respiration) and constituted the electrical double-layer capacitance of the carbon matrices^31^. Periodic release of the stored electrons through the bioelectrochemical circuit generated a set of discharging current (**SI Figure S4**). By integrating the discharging current with the incubation time, we quantified a total of 120±62 µmol e^-^ g^-1^ soil carbon that were snorkeled by the capacitive electron transfer. This number only accounted for 15% of the number of snorkeled electrons through the conductive electron transfer, which was probably due to the slower electron snorkeling kinetics of the capacitive electron transfer as expressed by the longer microbial adaptation phase and smaller current slope at the exponential growth phase (**Figure 1b**). Therefore, only at the late stage of the incubation did we show a decreased CH_4_ production rate (red line in **Figure 1a**). Significant amount of CH_4_ suppression is expected to occur only after an extended electron snorkeling period through capacitive electron transfer.

We investigated the effect of redox-cycling electron transfer on CH_4_ production in the microcosm peat-soil incubations (**SI Figure S3b**) for which the conductive and capacitive electron transfers were effectively eliminated (**SI Method S4**). Results showed that addition of pyrogenic carbon caused an instant drop in CH_4_ production and the total production (41±2.1 µmol g^-1^ soil carbon, orange and green lines in **Figure 1c**) was 13% lower than the pyrogenic carbon-free control treatment (47±2.5 µmol g^-1^ soil carbon, black line in **Figure 1c**). In accordance with the CH_4_ suppression, we observed an increased net accumulation of electrons (147±43 µmol e^-^ g^-1^ soil carbon) in the pyrogenic carbon-amended peat soil in comparison with the pyrogenic carbon-free control treatment (**Figure 1d** and **SI Figure S5**). This net electron accumulation was most likely derived from the alternative respiration that was enhanced by additional electron acceptance of the quinone groups on pyrogenic carbon. After accepting electrons, the quinone groups were reduced to phenol groups, which subsequently donated the accepted electrons (at least partly) to the soil organic matter and increased the electron accumulation in the peat soil. Such repeated redox-cycling electron transfer completed the electron snorkeling process and reduced CH_4_ production.

Isotopic analyses showed that during the redox-cycling electron transfer, a portion of the CO_2_ and CH_4_ production was derived from the metabolism of pyrogenic carbon (**Figure 2a, SI Method S5**, and **Table S1** and **S2**). The easily mineralizable carbon phase (characterized by volatile matter (35-23% w/w) and water extractable carbon (348-211 mg kg^-1^)) is enriched in the pyrogenic carbon that was produced at low (400-500°C) pyrolysis temperatures^32^, which was most likely the available carbon source for microbial metabolization^33^. Although the pyrogenic carbon-derived gas production only accounted for less than 2% of the total gas production, the metabolism of pyrogenic carbon could induce a shift of the peat-soil microbial structure and function^34^, which contributed to the suppression of the methanogenesis activity in addition to the redox-cycling induced electron snorkeling process. However, no detectable metabolism of pyrogenic carbon (**Figure 2b**) was observed during the conductive and capacitive electron transfers due to the lack of easily mineralizable carbon phase in the pyrogenic carbon that was produced at high (800°C) pyrolysis temperature^32^.

**Figure 2.**
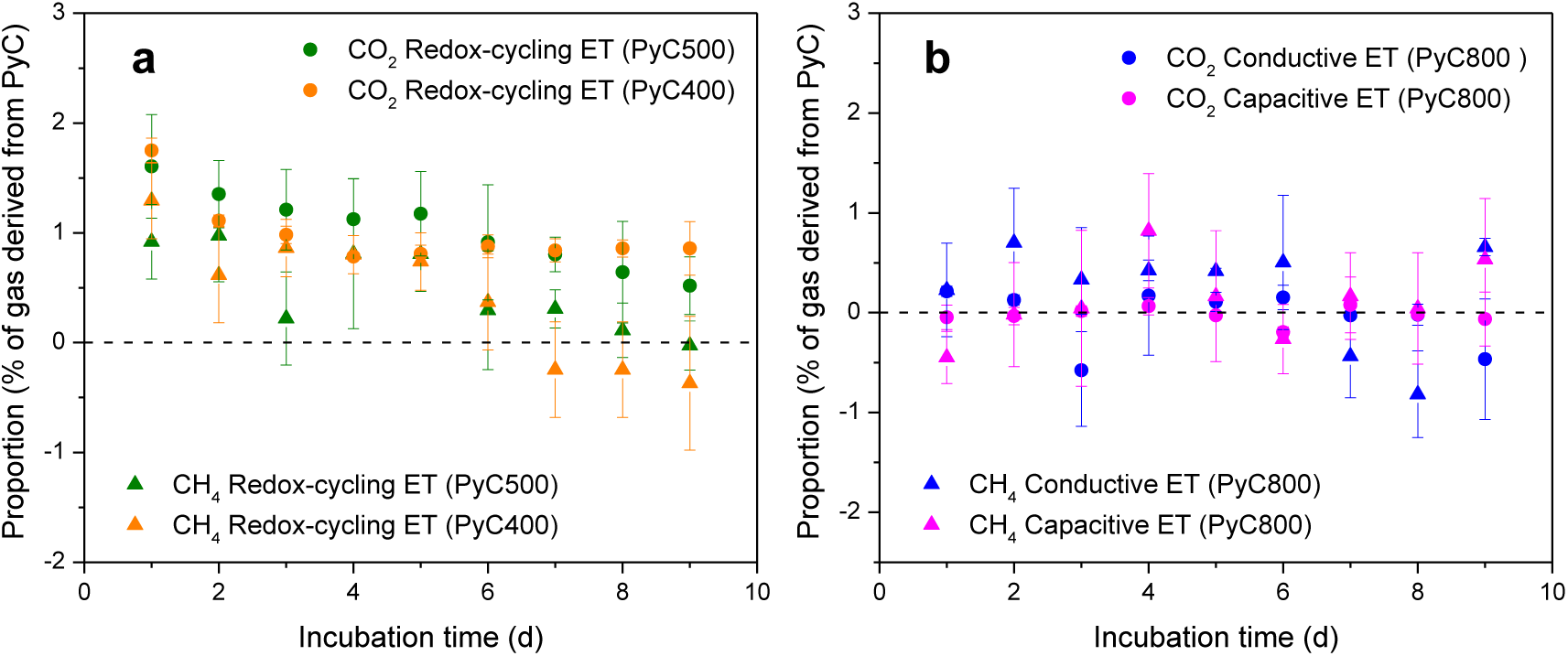
Proportions of CO_2_ and CH_4_ that were derived from the metabolism of pyrogenic carbon. **a.** Pyrogenic carbon-derived CO_2_ and CH_4_ production during the redox-cycling electron transfer (ET) of the pyrogenic carbon functional groups in the microcosm peat-soil incubations. Pyrogenic carbon was produced at 400 and 500°C (PyC400 and PyC500) and the application rate was 3 mg pyrogenic carbon g^-1^ soil. **b.** Pyrogenic carbon-derived CO_2_ and CH_4_ production during the conductive and capacitive ET through the pyrogenic carbon matrices in the bioelectrochemical peat-soil incubations. Pyrogenic carbon was produced at 800°C (PyC800) and the application rate was 10 mg pyrogenic carbon g^-1^ soil. Dashed lines in **a** and **b** indicate the proportions generated from the pyrogenic carbon-free control treatments. Calculation of proportion can be found in **SI Method S5** and **Table S1 and S2**. Day 0 in the x axes in **a** and **b** indicate the time of adding pyrogenic carbon into the peat soil.

### Electron snorkeling kinetics as a function of pyrolysis temperatures

Natural fires generate a highly heterogeneous pyrolysis temperature, which normally covers a range from 400 to 800°C depending on the size and duration of the fires^35,36^. In some extreme conditions, such as for the burning of wood core, or peat burning, the temperature can exceed 1000°C^37,38^. We investigated the determination of the pyrolysis temperature on the electron snorkeling kinetics by using the bioelectrochemical and microcosm pure-culture (*Geobacter sulfurreducens* strain PCA) incubations (**SI Method S6** and **S7** and **Figure S3c** and **d**). Compared to the multi-phase composition and complex microbiota in the peat-soil incubations, the pure-culture incubations provided a highly defined system for evaluating the kinetics across a single and identical microbe-pyrogenic carbon interface.

Both conductive and capacitive electron transfer mechanisms showed faster electron snorkeling kinetics (demonstrated by higher exponential current slopes in **Figure 3a** and **b**) through the carbon matrices that were produced at high temperature range (700-800°C) than low to intermediate temperature range (400-650°C). This result was in consistence with previous abiotic tests that the carbon matrices produced at high temperature range possesses more ordered polyaromatic carbon ring structures^24^, and thus higher electrical conductivity and capacitance^16^. In contrast to the conductive electron transfer, the capacitive electron transfer demonstrated slower electron snorkeling kinetics (**SI Figure S6-S9**), even though they share the same carbon structure in the carbon matrices to snorkel electrons. A similar phenomenon was also observed in the peat-soil incubations. We attributed this slower kinetics for the capacitive electron transfer to the build-up of an overpotential in the carbon matrices during the electron storage period, which slowed down further electron snorkeling until an electron release period was performed.

**Figure 3.**
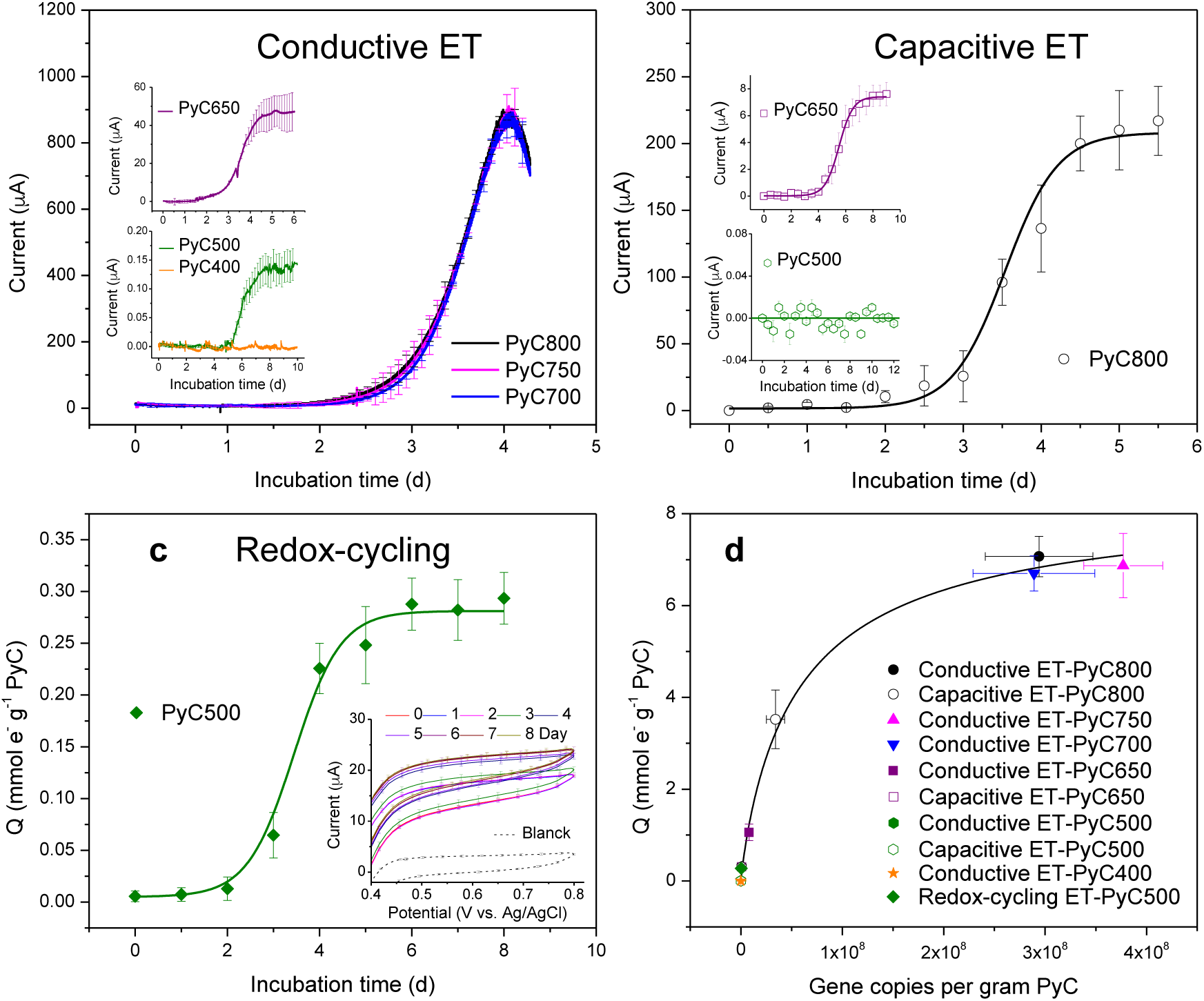
Electron snorkeling kinetics as a function of pyrolysis temperatures determined in the pure-culture (*G. sulfurreducens*) incubations. **a.** Current profiles generated from the conductive electron transfer (ET) through the pyrogenic carbon matrices. **b.** Discharging current generated from the capacitive ET through the pyrogenic carbon matrices. Both conductive and capacitive ET were tested in the bioelectrochemical pure-culture incubations. Pyrogenic carbon was produced at 400-800°C (PyC400-PyC800). The intermittent current signals in the capacitive ET showed the highest current point of each discharging period at a 0.1 s^-1^ recording frequency. Full discharging current profile could be found at the chronoamperograms in **SI Figure S7-S9. c.** Number of snorkeled electrons (Q) during the redox-cycling ET of the pyrogenic carbon functional groups. These numbers were background corrected, therefore represented the net number of snorkeled electrons (**SI Method S7**). Redox-cycling ET was tested in the microcosm pure-culture incubations. Pyrogenic carbon was produced at 500°C (PyC500). Inset: the oxidation current of ferrocyanide after reacting with pyrogenic carbon. Day 0 in the x axes in **a**-**c** indicate the time of inoculation of *G. sulfurreducens*. **d.** The increased number of snorkeled electrons as a function of the increased copy numbers of 16S rRNA genes of *G. sulfurreducens* (**SI Figure S13**) that was respiring on pyrogenic carbon by using different electron transfer mechanisms.

We determined the electron snorkeling rate of redox-cycling electron transfer (pyrogenic carbon produced at 500°C) at 0.16±0.02 mmol e^-^ g^-1^ pyrogenic carbon day^-1^, based on the increased number of snorkeled electrons at the exponential phase (**Figure 3c, SI Method S7**, and **SI Figure S10**). This rate was similar to the snorkeling rate of the conductive electron transfer through the carbon matrices produced at intermediate pyrolysis temperature 650°C (0.14±0.03 mmol e^-^ g^-1^ pyrogenic carbon day^-1^), but slower than that of the conductive and capacitive (3.22±0.15 and 0.54±0.08 mmol e^-^ g^-1^ pyrogenic carbon day^-1^, respectively) electron transfer through the carbon matrices produced at high pyrolysis temperatures (700-800°C). This trend indicated that while the conductive and capacitive electron transfer dominated the electron snorkeling process of the pyrogenic carbon that was produced at high pyrolysis temperatures, the redox-cycling electron transfer became more kinetically preferred in snorkeling electrons for the pyrogenic carbon that was produced at low to intermediate pyrolysis temperatures (500-650°C).

Even though the conductive electron transfer through the carbon matrices that were produced at high temperature range showed a rapid electron snorkeling kinetics, its overall performance was highly dependent on the available terminal electron acceptors. We found that dropping the potential gradient by 0.3 V, which is roughly equivalent to the shifting of the terminal electron acceptor from nitrate to iron minerals^11,39^, caused a 90% (N=3, P<0.01) decrease of the electron snorkeling kinetics (**Figure 4** and **SI Figure S11**). In contrast, the capacitive and redox-cycling electron transfers were less dependent on terminal electron acceptors due to their respective self-electron storage and acceptance functions. These functions enable pyrogenic carbon to act as an time-uncoupled intermediate electron acceptor, which sustains the electron snorkeling during times when a temporary lack of attached terminal electron acceptors exist^40^. The capacity of the self-electron storage and acceptance can be restored by releasing (**Figures 3b**) and donating (**Figures 3c**) electrons to available terminal electron acceptors when environmental conditions change. Such condition change can be triggered by, for example, oxygen penetration into temporary peat soils^11,41^ or the iron aggregation with pyrogenic carbon^42,43^. With the increased exposure frequency and duration to terminal electron acceptors, we expect that the electron snorkeling induced by capacitive and redox-cycling electron transfer will be more and more continuous similar to that induced by conductive electron transfer.

**Figure 4.**
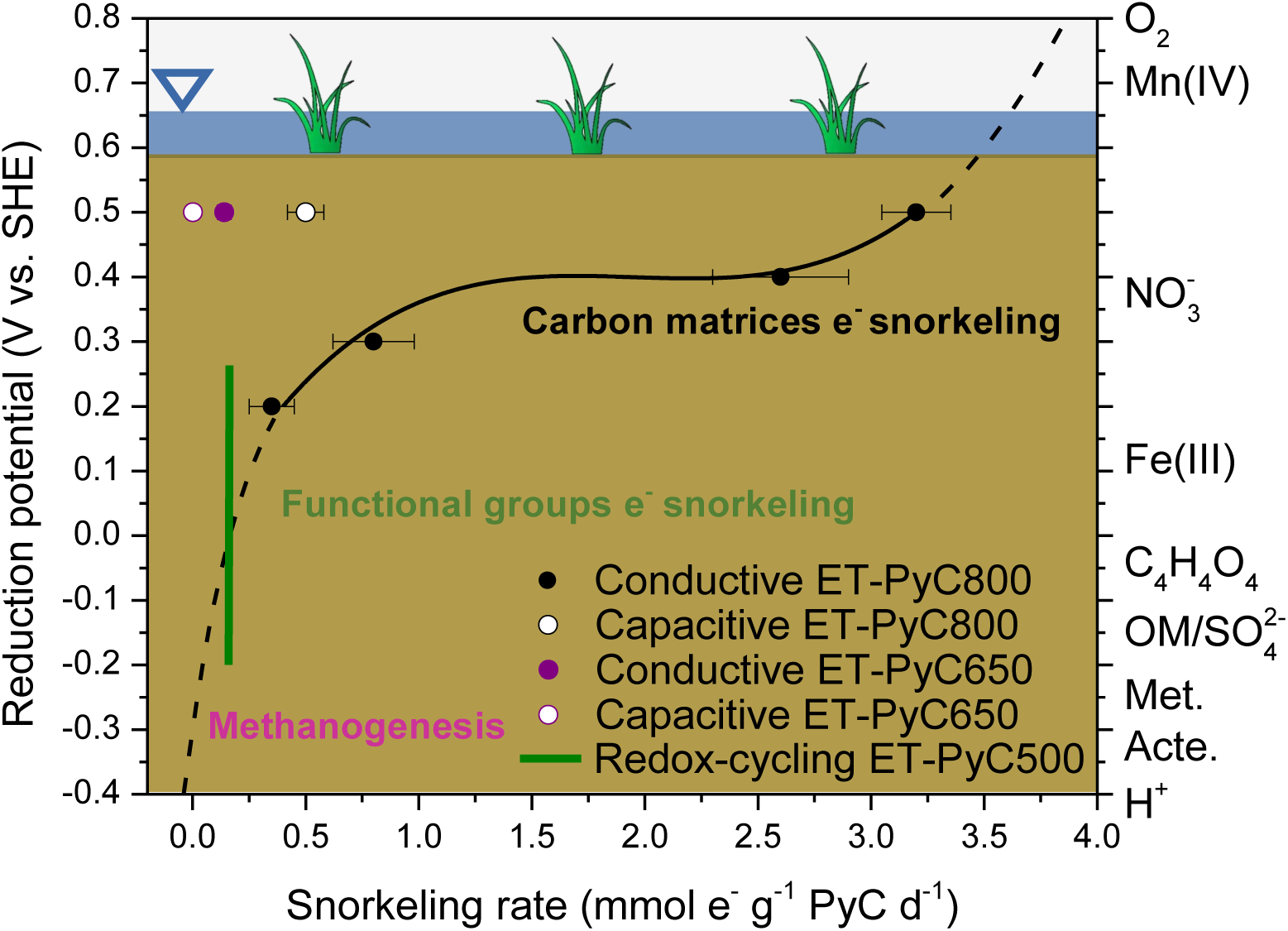
Dependency of electron snorkeling kinetics on environmental terminal electron acceptors. The solid black line is the trend line of measured data and the dashed black line is the extrapolated trend. The reduction potentials and the corresponding terminal electron acceptors listed on the y axes were adopted from ref^11,39,48^. The reduction potential (green line) of pyrogenic carbon functional groups was estimated based on direct electrochemical measurement^16^ and mediated iron^49^ and nitrate^50^ reduction. In the figure legend, ET stands for electron transfer and PyC and the following numbers indicate pyrogenic carbon and pyrolysis temperatures, respectively.

### Re-evaluating the legacy effect of pyrogenic carbon

Here, we presented a post-fire CH_4_ suppression phenomenon induced by fire-derived pyrogenic carbon in a northern peat soil. In contrast to the traditional ideas that rationalize the legacy effect of pyrogenic carbon mainly from the perspective of its persistence^44,45^, the CH_4_ suppression observed in this study was a result of strong bioelectrochemical and redox interactions between pyrogenic carbon and microbial respiration. The CH_4_ was suppressed by the electron snorkeling process of pyrogenic carbon, which redirected the electron fluxes towards the organic matter in peat soil as the terminal electron acceptor to facilitate alternative respiration by *Geobacter* spp.. Both pyrogenic carbon matrices and functional groups contributed to electron snorkeling by employing the conductive, capacitive, and redox-cycling electron transfer mechanisms. This multi-electron transfer mechanism allowed pyrogenic carbon to process the electron snorkeling across a wide range of naturally occurring fire/pyrolysis temperatures (**Figure 3a-c**). The 13-24 % reduction in CH_4_ production by pyrogenic carbon electron snorkeling shown in this study translates into average global reduction of 35 Tg CO_2_e annually, equivalent to the greenhouse gas emissions of 7,600,000 cars (**SI Method S8**). Therefore, CH_4_ suppression by pyrogenic carbon electron snorkeling could be responsible for a considerable effect in neutralizing the negative climate impact of forest fires. However, many further studies would be necessary to confirm this.

Increased accumulation of pyrogenic carbon is expected as a result of more frequent forest fires^46^. A higher amount of pyrogenic carbon caused greater activity of alternative respiration (**SI Figure S12**). However, the number of snorkeled electrons did not show an accelerated increase in response to the increased biomass in alternative respiration. Instead, the increasing rate decelerated and eventually leveled out (**Figure 3d** and **SI Figure S13**). This response pattern resembles the Michaelis–Menten behavior in the enzymatic chemistry, which reflects the catalytic nature of the electron snorkeling process. On the other hand, the decelerated increasing rate implies a functioning ceiling existed in the electron snorkeling process, which limits further CH_4_ suppression even though the input of pyrogenic carbon is sufficient. Therefore, we suggest future studies should focus on investigating to what extent pyrogenic carbon can suppress peat CH_4_ production and on quantifying how much contribution this suppression can make to offset the negative climate impact caused by fire emission. Such studies will potentially lead to a paradigm shift of the pyrogenic carbon legacy effect from passive carbon sequestration to active mitigation of greenhouse gas emissions from the host soil.

## Methods

### Pyrogenic carbon samples

Pyrogenic carbon samples were produced under controlled conditions in the laboratory by anoxic pyrolysis of woody biomass (shrub willow) with pyrolysis temperatures of 400-800°C, which covered the temperature range of naturally occurring forest and grassland fires^35,36^. We used a dwell time of 60 min for all pyrolysis temperatures. Before pyrolyzing, the biomass carbon had been labelled in growth chambers using ^13^C labeled CO_2_ as the source for willow growth. The bulk δ^13^C in pyrogenic carbon after biomass pyrolysis was on average 774±2.3‰ (vs. VPDB). More information on the physicochemical properties (carbon content, elemental composition, pH, etc) of pyrogenic carbon can be found in our previous study^32^.

### Peat soil sample

We sampled the peat soil from the Mclean Bog located in Dryden, New York (42°30’ N, 76°30’ W). The Mclean Bog is a national natural landmark with known methanogenesis activity and CH_4_ production. Soil was collected from depths that were approximately 10-15 cm below the surface and contained 52% total carbon and 2.1% total nitrogen (dry weight percentage). Detailed description of the peat soil was given in **SI Method S1**. For all peat-soil incubations, 10 g wet soil was suspended with deoxygenated and deionized water to reach a final volume of 30 ml. The final pH of the soil suspension was 4.5, and all pyrogenic carbon had been adjusted to this pH prior to incubation.

### Pure-culture inoculant

The bacterial pure-culture *G. sulfurreducen*s strain PCA (1 mL of stock culture at 0.1 OD) was inoculated in all pure-culture incubations. From all electron donating bacteria, we chose *G. sulfurreducens* because of its mediator-free electron transfer mechanisms that facilitated the mechanism study by only crossing one electron transfer interface (i.e., bacteria-pyrogenic carbon interface instead of bacteria-mediator-pyrogenic carbon interfaces) and avoided potential interfere caused by the mass transport of mediators. Further, *G. sulfurreducens* is an important and abundant alternative-respiring bacterium that coexists with methanogens in many anaerobic environments^47^. The average abundance of *Geobacter* species accounted to 1% of 16S rRNA gene sequences, which dominated the alternative respiratory bacteria in the studied peat soil (**SI Figure S2**).

### Bioelectrochemical and microcosm incubations

The bioelectrochemical and microcosm incubations were performed to investigate the electron snorkeling induced by the pyrogenic carbon matrices and functional groups, respectively. We performed peat-soil and pure-culture incubations in parallel together with pyrogenic carbon in bioelectrochemical and microcosm incubations. The number of snorkeled electrons in the bioelectrochemical incubations was quantified based on the current signal induced by either conductive or capacitive electron transfer through the bioelectrochemical circuit. The number of snorkeled electrons in the microcosm incubations was determined by the electron accumulation due to the redox-cycling electron transfer in the peat soil or growth medium.

Application rate was 0 and 10 mg pyrogenic carbon g^-1^ soil and 0 and 6.7 mg pyrogenic carbon mL^-1^ growth medium for bioelectrochemical peat-soil and pure-culture incubations, respectively. Application rate was 0, 0.03 and 3 mg pyrogenic carbon g^-1^ soil in the microcosm peat-soil incubations and 0 and 1 mg pyrogenic carbon mL^-1^ growth medium in the microcosm pure-culture incubations. We monitored daily production rates of CO_2_ and CH_4_ and δ^13^C partitioning in the gas phase of all bioelectrochemical and microcosm incubations, using a Picarro stable isotope analyzer (G2201-I, Santa Clara, CA, USA). Detailed information about the design, operation and data acquisition of bioelectrochemical and microcosm incubations could be found in **SI Method S2-S7**.

## Acknowledgements

This research was supported by NSF-BREAD (grant number IOS-0965336), USDA NIFA Carbon Cycles (2014-67003-22069). The authors acknowledge support from the Alexander von Humboldt Foundation in the framework of the Alexander von Humboldt Professorship endowed by the Federal Ministry of Education and Research in Germany. Any opinions, findings and conclusions or recommendations expressed in this material are those of the authors and do not necessarily reflect the views of the donors.

## Author contributions

T.S., J.L., and L.T.A. planned the research, T.S. conducted the bioelectrochemical and microcosm incubation experiments, laboratory analysis, and data analysis. T.S. wrote the manuscript in close collaboration with L.T.A. J.J.L.G. assisted with the bioelectrochemical measurements and interpretation and qPCR analysis. J.D.S. and J.B.Y. provided the peat-soil sample and performed the physicochemical and genomic analyses. A.E. assisted the setup of incubation and gas and isotopic analysis. All authors edited the manuscript.

## Additional information

Supplementary information as mentioned in the manuscript is available online.

## Competing interests

The authors declare no competing financial interests.

