## Supplementary Information for "Suppressing peatland methane production by electron snorkeling through pyrogenic carbon"

### for

Number of pages: 30
Number of methods: 8
Number of figures: 13
Number of tables: 2

### Supplementary Methods

#### Method S1: Peat soil samples

The peat soil samples were collected in June at the Mclean Bog located in Dryden, New York (42°30' N, 76°30' W). Soils were sampled at several locations of the center of the bog, from depths approximately 10-15 cm below the water level. McLean Bog is an ombrotrophic (= rain fed) peat site, which has an area of 0.004-km$^2$. Total peat depth is 8 m. *Sphagnum* species include *S. angustifolium* and *S. amgellanicum*. Shrubs include *C. calyculata* and *V. corymbosum* (highbush blueberry). The sedge *Eriophorum vaginatum* (cotton sedge) is also commonly distributed. Further descriptions can be found in our previous studies[1-3]. Gas emission profile from the peat soil was given in **Figure S1a-d**.

*Microbial Sequencing Analysis.* For community sequencing, DNA was extracted from peat samples with MoBio (now QIAGEN) Laboratories PowerSoil® DNA Isolation Kit and cleaned using the PowerClean® kit, following the manufacturers protocol with a heating step (65 °C for 30 minutes) added during the DNA extraction following bead beating. Sequences were collected on the MiSeq platform by the Department of Energy's (DOE) Joint Genome Institute (JGI) for bacteria and archaea using the V4 region of SSU rRNA (515/806) primer pairing. Raw sequence data was obtained from the JGI database and sequences were first quality filtered using BBMap package[4]. PANDAseq[5] was used to align forward and reverse reads, and aligned sequences were then processed with QIIME[6], and USEARCH version 8[7] software using a 97% confidence value for OTU assignment[8,9]. Contigs were generated from complementary forward and reverse sequences while discarding sequence reads shorter than 240 base pairs (bp) or longer than 350 bp. The Greengenes 2013 database[10] was used for taxonomic assignment of bacterial and archaeal assemblages. Taxonomic identification data was generated and visualized using QIIME 1 scripts[6]. Microbial community structure of the peat soil was given in **Figure S2**.

*Alternative terminal electron acceptors in the peat soil.* The average production ratio of $CO_2$ to $CH_4$ dropped from 15 to 8 during the 9 days of incubation period (**Figure S1e and f**), which was comparable to the ratios (range 1:1 to 40:1) reported by previous studies on several northern peat soils[3,11-13]. Higher $CO_2$ production than $CH_4$ indicated that alternative terminal electron acceptors (other than $O_2$) existed in the anaerobic peat soil. Iron is a common alternative electron acceptor found in peat soils and contributes to alternative respiration not only influencing the mitigation of methanogenesis[14,15] but also the anoxic methane oxidation[16]. The peat soil used in this study, however, was from an ombrotrophic site which contains no inlet or outlet streams so that the iron inputs are usually insignificant. The measured iron concentration in the peat porewater is 4.2±1.3 μmol $L^{-1}$ at 0-40 cm depth and 1.8±0.1 μmol $L^{-1}$ at 100 cm depth. It has also been reported that the iron only contributed less than 2% to the total electron exchange capacities of ombrotrophic peat soils[17]. In comparison, the high carbon content (52%, dry weight percentage) in the peat soil indicated that organic matter, particularly its quinone moieties[18], was the major alternative terminal electron acceptor. Organic matter has been widely studied and demonstrated its importance in accepting electrons in anoxic soil conditions[19,20], especially under long-term reducing conditions after the exhaustion of such as $NO_3^-$, iron and manganese phases, and $SO_4^{2-}$ ref[3]. The highest number of electron acceptance in the peat soil was determined at 785±120 μmol $e^-$ $g^{-1}$ soil carbon in the bioelectrochemical peat-soil incubations **(Figure 1b** in the main text), which was close to the reported electron accepting capacities of organic matters (a few hundred to thousand μmol $e^-$ $g^{-1}$ soil carbon) of several northern peat soils[17,21,22]. Another reason for the higher $CO_2$ production could be the introduction of oxygen into the peat soil. Even though we used nitrogen protection during the incubation preparation (**Method S2**), small amount of oxygen penetration into the peat soil was inevitable.

**Method S2: Peat-soil incubation preparation**

We used two layers of Ziplock bags to store and transport each pack of soil samples to prevent the oxygen penetration. The soils were stored in the lab at room temperature and dark environment until the incubation started. For the incubation, a certain amount of the soil was picked in the middle of each sampling pack, which was then homogeneously mixed to make one stock soil sample. The stock soil sample has a water content of 90% and a pH of 4.1. Afterwards, we divided the stock soil sample into three 10-g portions as the triplicates of one set of incubation. Each 10-g portion was individually placed into a 50 mL beaker and submerged with DI water to 30 mL for a full suspension. The beaker was pre-fixed in a 180 mL jar using hot glue. The jar was equipped with a gas-tight lid that had several rubber septa for gas sampling and pyrogenic carbon addition (**Figure S3a** and **b**). 50 mL DI water was added in the jar (surrounding the beaker to keep 100% moisture), which left the gas phase volume of each jar at 130 mL. The above-described preparation process was repeated for each individual incubation until fulfilling the designed incubation purposes. All preparations were performed in an anaerobic box with a continuous flow of $N_2$ gas. The closed jar was fully flushed once again with $N_2$ gas immediately prior the start of the incubation. A magnetic stir bar (400 rpm) was used in the soil suspension to maintain homogenous during the incubation.

**Method S3: Bioelectrochemical peat-soil incubation**

The effect of conductive and capacitive electron transfer of the pyrogenic carbon matrices on electron snorkeling was investigated in bioelectrochemical peat-soil incubations. A two-chamber bioelectrochemical system was employed (**Figure S3a)** in which a pyrogenic carbon rod was the working electrode (WE) and placed in the peat soil chamber along with a Ag/AgCl (saturated KCl) reference electrode (RE). The graphite rod counter electrode (CE) was placed in a separated chamber to avoid any generation of hydrogen in the peat soil chamber, which could potentially enhance the methanogenesis and methane emission. The bioelectrochemical peat-soil incubations were carried out with the soil native microbiota (i.e., without any further inoculation or nutrient addition) and no autoclavation was performed. The incubation temperature was maintained at 32°C in an incubation room. Lights were off all the time except for sampling. The gas phase was measured once a day by the Picarro stable isotope analyzer (G2201-I, Santa Clara, CA, USA). After each measurement, the gas phase of incubation was completely replaced by nitrogen gas.

Electrical potentials were applied at the pyrogenic carbon WE to provide sufficient driving force and facilitate the terminal electron acceptance. By eliminating the electron snorkeling limit induced by the terminal electron accepting step, we were able to resolve the controlling effect of only conductive or capacitive electron transfer on the overall electron snorkeling process. The potential was applied constantly at the pyrogenic carbon WE (+0.5 V vs. SHE, depicted by the red solid line in **Figure S3a**) to guarantee a continuous acceptance of the electrons that was snorkeled by the conductive electron transfer. In contrast, the potential was applied intermittently (+0.5 V vs. SHE, depicted by the red dash line in **Figure S3a**) to accept the electrons that were periodically snorkeled through a series of electron storage and release cycles in the capacitive electron transfer. The potential was applied once every 11.5 h during the capacitive electron transfer and each application period lasted for 0.5 h to ensure that the electrons were intermittently snorkeled only by the capacitive electron transfer other than the continuous electron snorkeling induced by the conductive electron transfer. Both conductive and capacitive electron transfers induced passing of electric current through the bioelectrochemical circuit **(Figure 1b** in the main text and **Figure S4**). By integrating the current as a function of incubation time, we quantified the number of snorkeled electrons in conductive and capacitive electron transfers.

### Method S4: Microcosm peat-soil incubation

The effect of redox-cycling electron transfer of the pyrogenic carbon functional groups on electron snorkeling was investigated in microcosm peat-soil incubations (**Figure S3b**). Pyrogenic carbon produced at 400-500°C was used to study the redox-cycling electron transfer, due to its enrichment of functional groups (oxygen:carbon and hydrogen:carbon ratios range from 0.15 to 0.11 and 0.6 to 0.4, respectively). In contrast, the oxygen:carbon and hydrogen:carbon ratios of the pyrogenic carbon produced at 800°C are only 0.06 and 0.14[23], which indicated a condensed carbon structure and lack of functional groups. Due to the enrichment of functional groups and less condensed carbon structure, the conductive and capacitive electron transfers through the carbon matrices that were produced at 400-500°C were highly diminished (green and orange lines in **Figure 3a** and **b** in the main text). Therefore, any electron snorkeling process occurred through the pyrogenic carbon that was produced in this low temperature range was a result of the redox-cycling electron transfer of the functional groups. The microcosm peat-soil incubations were carried out with the soil native microbiota (i.e., without any further inoculation or nutrient addition) and no autoclavation was performed. The incubation temperature was maintained at 32°C in an incubation room. Lights were off all the time except for sampling. The gas phase was measured once a day by the Picarro stable isotope analyzer (G2201-I, Santa Clara, CA, USA). After each measurement, the gas phase of incubation was completely replaced by nitrogen gas.

Pyrogenic carbon powder was added in the microcosm peat-soil incubations and without any application of electrical potentials. Therefore, the driving force of the redox-cycling electron transfer was dependent on the inherent reduction potential of the functional groups (-0.2 to +0.25 V for quinone/phenol functional groups[24-27]), which snorkeled electrons by spontaneous electron accepting (from alternative respiration) and donating (to alternative terminal electron acceptor) cycles. The number of snorkeled electrons by the redox-cycling electron transfer was quantified by the electron accumulation in the peat soil (i.e., the electrons that were accepted by alternative terminal electron acceptor and hold in the peat soil other than lost in the form of $CH_4$ emission). Electron accumulation was quantified by the increased electron donating capacity of the peat soil, using a previously reported hydrodynamic cyclic voltammetric method[28]. Briefly, 1 mL soil suspension was sampled out at day 1, 3, and 9 of each treatment and mixed with a 54 mL ferricyanide solution. This total 55 mL mixture contained 0.36 g dry soil carbon $L^{-1}$ solution, 10 mM potassium ferricyanide, and 3 M NaCl as the supporting electrolyte. After overnight shaking and reacting, a cyclic voltammetry was performed in the mixture using a glassy-carbon rotating-disk electrode (6 mm diameter). Due to the reduction of ferricyanide and production of ferrocyanide, an increased oxidation current ($j_{Ferro}$, A $cm^{-2}$) appeared in the hydrodynamic cyclic voltammograms (**Figure S5**). The electron donating capacity of the peat soil was quantified by measuring the final production of ferrocyanide using eq. S1 and S2:

$$Q_{ED} = \frac{nV[\text{Ferro}]}{m} \qquad \text{eq. S1}$$

$$[\text{Ferro}] = \frac{j_{\text{Ferro}}}{0.62nFD_{Ferro}^{2/3}v^{-1/6}\omega^{1/2}} \qquad \text{eq. S2}$$

in which $Q_{ED}$ is the number of donated electrons from the peat soil (mol $e^{-}$ $g^{-1}$ soil carbon), which was projected to the number of accumulated electrons in the peat soil, n = 1 is the number of electrons exchanged per mol ferricyanide reduction, V is the volume of solution (55 $cm^3$), m is the mass of dry soil carbon (0.02 g), and the [Ferro] is the ferrocyanide concentration (mol $cm^{-3}$). F is the Faraday constant (96,485 C $mol^{-1}$), $D_{Ferro}$ is the diffusion coefficient of ferrocyanide at 30°C, $4.27 \times 10^{-6}$ $cm^2$ $s^{-1}$ ref[28], ν is the kinematic viscosity of 3 M NaCl solution at 30 °C ($9.83 \times 10^{-3}$ $cm^2$ $s^{-1}$, ref[28]), ω is the RDE rotation speed (104.7 rad $s^{-1}$).

**Method S5: Calculation of the proportion of gases emitted from the respiration of pyrogenic carbon**

In all peat-soil incubations, the added pyrogenic carbon was isotopically labelled with $^{13}C$, which resulted in a $\delta^{13}C$ of pyrogenic carbon at 774±2.3‰. We monitored the $\delta^{13}CO_2$ and $\delta^{13}CH_4$ production during the bioelectrochemical (**Table S1**) and microcosm (**Table S2**) peat-soil incubations to track the respiration of pyrogenic carbon by the native peat-soil microbiota. The percentage of $CO_2$ and $CH_4$ (relative to the total gas emission from the peat soil) that were derived from the respiration of pyrogenic carbon were calculated based on eq. S3[23]:

$$P_{CO_2\ or\ CH_4} = \frac{\delta^{13}C_{PyC+soil} - \delta^{13}C_{soil}}{\delta^{13}C_{PyC} - \delta^{13}C_{soil}} \times 100 \qquad \text{eq. S3}$$

in which $P_{CO_2\ or\ CH_4}$ is the percentage of $CO_2$ or $CH_4$ that are derived from the respiration of pyrogenic carbon in the bioelectrochemical and microcosm peat-soil incubations, $\delta^{13}C_{PyC+soil}$ represents the $\delta^{13}C$ values of the peat-soil incubations with pyrogenic carbon, $\delta^{13}C_{soil}$ is the $\delta^{13}C$ values of the pyrogenic carbon-free control incubations (i.e., the peat soil only), $\delta^{13}C_{PyC}$ is the $\delta^{13}C$ value of the labelled pyrogenic carbon. The $\delta^{13}C$ values in terms "$\delta^{13}C_{PyC+soil}$" and "$\delta^{13}C_{soil}$" are either $\delta^{13}C$ of $CO_2$ or $CH_4$ as given in **Table S1** and **S2**.

**Method S6: Bioelectrochemical pure-culture incubation**

We investigated the determination of the pyrolysis temperature on the electron snorkeling kinetics of conductive and capacitive electron transfers using the bioelectrochemical pure-culture incubations. A one-chamber bioelectrochemical system (**Figure S3c**), adapted from previously published studies[29,30], was used in the bioelectrochemical pure-culture incubations due to its autoclavable and easy-assemble features. All pyrogenic carbon WE, Ag/AgCl RE, and graphite CE electrodes were placed in the same chamber. The bioelectrochemical systems were autoclaved, filled with 15 mL of sterile growth media (consisted of 2.5 g $NaHCO_3$, 0.25 g $NH_4Cl$, 0.52 g $NaH_2PO_4$, 1 g KCl, 1 mL of vitamin mix, 1 mL of mineral mix, and 40 mM sodium acetate per litter), and placed in a 30°C water bath. $N_2$:$CO_2$ (80%:20%) gas was continuously sparged in the growth media to maintain anaerobic condition. *G. sulfurreducen*s strain PCA (1 mL of stock culture at 0.1 OD) was inoculated as an alternative respiratory bacterium in all pure-culture incubations. Electrical potentials (+0.2 to +0.5 V vs. SHE) were applied either constantly (depicted by the red solid line in **Figure S3c**) or intermittently (depicted by the red dash line in **Figure S3c**) at the pyrogenic carbon to eliminate the electron snorkeling limit induced by the terminal electron accepting step. Therefore, the overall electron snorkeling kinetics was only controlled by either conductive or capacitive electron transfer of carbon matrices. Pyrogenic carbon that was produced from low to high pyrolysis temperatures (400-800°C) was incubated in the bioelectrochemical pure-culture incubations. The potential was applied once every 11.5 h during the capacitive electron transfer and each application period lasted for 0.5 h to ensure that the electrons were intermittently snorkeled only by the capacitive electron transfer other than the continuous electron snorkeling induced by the conductive electron transfer. Both conductive and capacitive electron transfers induced passing of electric current through the bioelectrochemical circuit (**Figure 3a and b** in the main text and **Figure S7-S9**). By integrating the current as a function of incubation time, we quantified the number of snorkeled electrons in conductive and capacitive electron transfers.

**Method S7: Microcosm pure-culture incubation**

The microcosm pure-culture incubations were carried out in serum bottles with pyrogenic carbon powder added to the growth media and without any application to electrical potentials (**Figure S3d**). 50 mL growth media was added into each serum bottle with *G. sulfurreducen*s strain PCA as an alternative respiratory bacterium. The composition of inoculum and growth media of *G. sulfurreducen*s was the same as described above in the bioelectrochemical pure-culture incubations (see **Method S6**). Pyrogenic carbon produced at 500°C was used to study the redox-cycling electron transfer in the microcosm pure-culture incubations. Due to the enrichment of functional groups and less condensed carbon structure, the conductive and capacitive electron transfers through the carbon matrices that were produced at 500°C were highly diminished (green lines in **Figure 3a** and **b** in the main text). Therefore, any electron snorkeling process occurred through the pyrogenic carbon that was produced in this low temperature was a result of the redox-cycling electron transfer of the functional groups. The addition rate of pyrogenic carbon was 1 mg pyrogenic carbon $mL^{-1}$ growth media.

During the incubation, pyrogenic carbon became more and more reduced due to the continuous accepting of electrons generated from *G. sulfurreducen*s respiration (i.e., the microbial reduction in eq. S4). Aliquots of the reduced pyrogenic carbon were sampled daily and re-oxidized by donating electrons to potassium ferricyanide (eq. S5).

$$\text{Acetate} + \text{PyC}_{\text{Ox}} \xrightarrow[\text{reduction}]{\text{Microbial}} \text{CO}_2 + \text{H}^+ + \text{PyC}_{\text{Red}} \qquad e^- \textit{ acceptance} \qquad \text{eq. S4}$$

$$\text{PyC}_{\text{Red}} + \text{Ferri} \xrightarrow[\text{oxidation}]{\text{Chemical}} \text{PyC}_{\text{Ox}} + \text{Ferro} \qquad e^- \textit{ donation} \qquad \text{eq. S5}$$

We determined the number of snorkeled electrons (Q, mol $e^-$ $g^{-1}$ pyrogenic carbon) of the complete redox cycle (eq. S3 + eq. S4) of pyrogenic carbon functional groups by measuring the final production of ferrocyanide (eq. S6[28]):

$$Q = \frac{nV[\text{Ferro}]_{\text{redox-cycling}}}{m} \qquad \text{eq. S6}$$

in which n = 1 is the number of electrons exchanged per mol ferricyanide reduction, V is the volume of solution (55 $cm^3$), m is the mass of pyrogenic carbon (0.01 g), and the $[Ferro]_{redox\ cycling}$ is the ferrocyanide concentration (mol $cm^{-3}$) produced by the redox cycling of the functional groups.

The $[Ferro]_{redox\ cycling}$ was measured by mixing 10 mL incubation solution with 45 mL potassium ferricyanide solution. This total 55 mL mixture contained 0.01 g pyrogenic carbon, 10 mM ferricyanide, and 3 M NaCl as the supporting electrolyte. After overnight shaking and reacting, a cyclic voltammetry was performed in the mixture using a glassy-carbon rotating-disk electrode (6 mm diameter). Due to the reduction of ferricyanide and production of ferrocyanide, an increased oxidation current ($j_{total}$, A $cm^{-2}$) appeared in the hydrodynamic cyclic voltammograms (**Figure S10**). After subtracting the background reduction induced oxidation current, we obtained the $[Ferro]_{redox\ cycling}$ based on the Levich equation (eq. S7):

$$[Ferro]_{redox\ cycling} = \frac{j_{total} - j_{inoculation} - j_{HQ}}{0.62 nFD_{Ferro}^{2/3} v^{-1/6} \omega^{1/2}} \quad \text{eq. S7}$$

where F is the Faraday constant (96,485 C $mol^{-1}$), $D_{Ferro}$ is the diffusion coefficient of ferrocyanide at 30°C, $4.27 \times 10^{-6}$ $cm^2$ $s^{-1}$ ref[28], ν is the kinematic viscosity of 3 M NaCl solution at 30 °C ($9.83 \times 10^{-3}$ $cm^2$ $s^{-1}$, ref[28]), ω is the RDE rotation speed (104.7 rad $s^{-1}$).

$j_{total}$ in eq. S7 indicates the total oxidation current (**Figure S10c**), $j_{inoculation}$ (**Figure S10b**) and $j_{HQ}$ (**Figure S10a**) in eq. S7 are the oxidation current induced by the background reduction. The background reduction derived from: (1) the immediate reduction of ferricyanide by the biofilm electrons after the inculation of *G. sulfurreducen*s (i.e., $j_{inoculation}$ in eq. S7); and (2) the abiotic reduction of ferricyanide by the inherent hydroquinone/phenol groups in pyrogenic carbon (i.e., $j_{HQ}$ in eq. S7). The $j_{inoculation}$ was assessed by inoculating microbes into the growth medium that contained sand but no pyrogenic carbon and other electron acceptors. The $j_{HQ}$ was determined in the growth medium with the addition of pyrogenic carbon but without microbe inoculation. By subtracting $j_{inoculation}$ and $j_{HQ}$ from $j_{total}$, we obtained the oxidation current that was only induced by the redox-cycling electron transfer of pyrogenic carbon in supporting the growth of *G. sulfurreducen*s.

**Method S8: Estimation of annual methane suppression by accumulation of pyrogenic carbon in northern peatlands**

Annually, peatland fires approximately produce 9 Tg pyrogenic carbon[31], which is averaged to a 4,000,000 $km^2$ peatland area[32,33] and yields a production rate of 2,250,000 g pyrogenic carbon $km^{-2}$ $yr^{-1}$ or 0.0062 g pyrogenic carbon $m^{-2}$ $d^{-1}$. The suppressed $CH_4$ by the electron snorkeling effect of unit gram of pyrogenic carbon is 1.3 mg (averaged from all electron transfer mechanisms, see **Figure 1** in the main text), which is divided by 9 days of incubation time and 0.001256 $m^2$ incubation area and gives the $CH_4$ suppressing rate at 114 mg $CH_4$ $m^{-2}$ $d^{-1}$ $g^{-1}$ pyrogenic carbon. Multiplying the production rate of pyrogenic carbon and its CH4 suppressing rate yields a total suppression of 0.7 mg $CH_4$ $m^{-2}$ $d^{-1}$ (i.e., 0.0062 g pyrogenic carbon $m^{-2}$ $d^{-1}$ × 1 $m^2$ × 1 d × 114 mg $CH_4$ $m^{-2}$ $d^{-1}$ $g^{-1}$ pyrogenic carbon). This number of suppressed $CH_4$ accounts for 1-14% of the total $CH_4$ production (5-80 mg $CH_4$ $m^{-2}$ $d^{-1}$)[34,35] in northern peatlands. Since $CH_4$ has a 34 times[36] stronger global warming potential than $CO_2$, the suppressed $CH_4$ is equivalent to a reduction of 24 mg $CO_2e$ $m^{-2}$ $d^{-1}$, or 35 Tg $CO_2e$ $yr^{-1}$ (i.e., 24 mg $CO_2e$ $m^{-2}$ $d^{-1}$ × 1,000,000 $m^2$ $km^{-2}$ × 4,000,000 $km^2$ × 365 d $yr^{-1}$). The average annual $CO_2$ emission of a passenger vehicle is 4,600,000 g (USEPA-420-F-18-008), therefore, 35 Tg $CO_2e$ $yr^{-1}$ is equivalent to the greenhouse gas emissions of 7,600,000 cars. Although simple, this calculation highlights the potentially large impacts of pyrogenic carbon induced electron snorkeling in neutralizing the negative climate impact of forest fires.

### Supplementary Figures

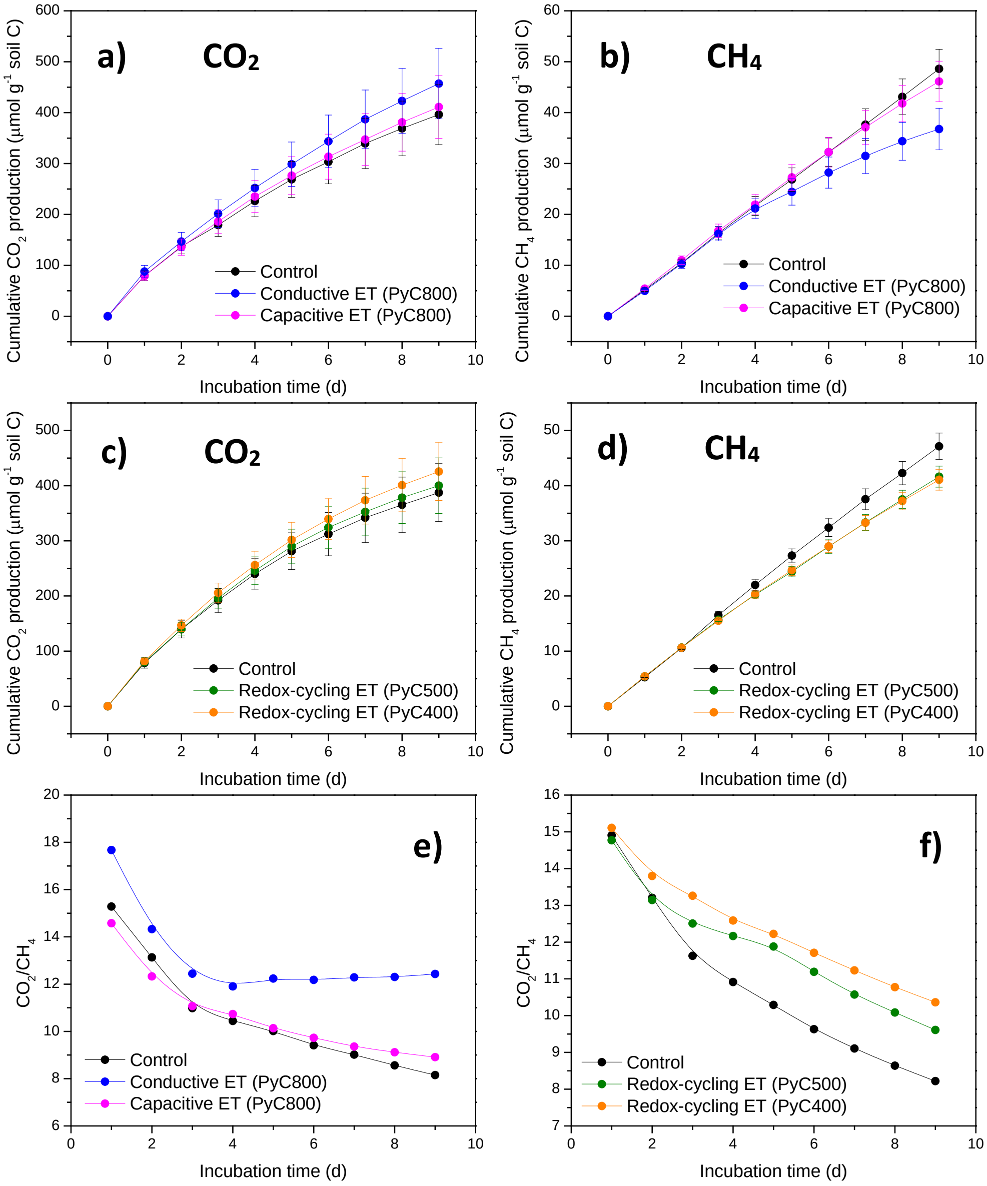


**Figure S1 | Total gas emission from the peat soil as a function of incubation time. a** and **b.** Total $CO_2$ and $CH_4$ emissions from the pyrogenic carbon-free control treatment (i.e., the original peat soil emission) and from the incubations that were associated with conductive and capacitive electron transfers (ET) for electron snorkeling in the bioelectrochemical peat-soil incubations. **c** and **d.** Total $CO_2$ and $CH_4$ emissions from the pyrogenic carbon-free control treatment and from the incubations that were associated with the redox-cycling ET for electron snorkeling in the microcosm peat-soil incubations. **e** and **f.** $CO_2$ to $CH_4$ ratios in the pyrogenic carbon-free control treatment and bioelectrochemical and microcosm peat-soil incubations. In all charts, PyC stands for pyrogenic carbon and the following numbers indicate the pyrolysis temperatures.

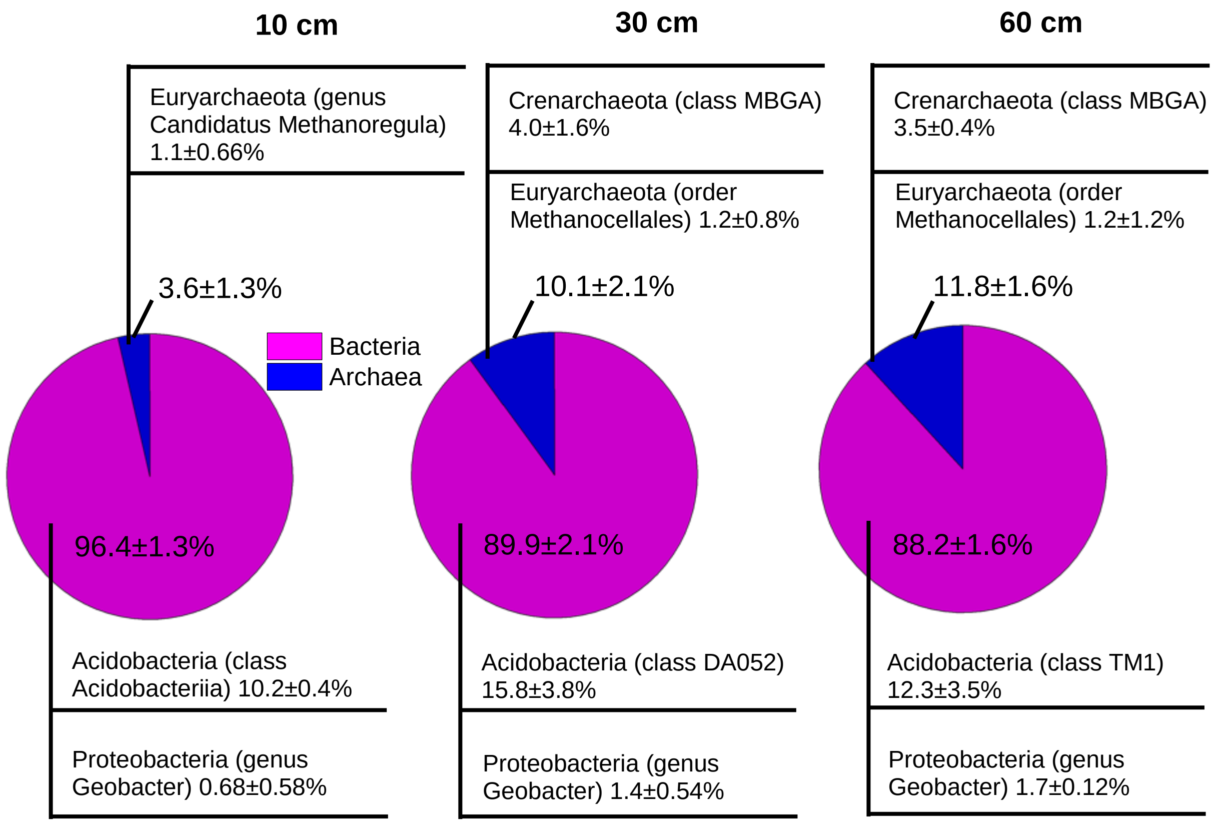


**Figure S2 | Microbial community analysis of the studied peat soil at different soil depth (10-60 cm).** We sampled the soil at 10-15 cm. The pie charts show the relative abundance of bacteria and archaea. The notes above the charts give the relative abundance of the most abundant archaea and methanogens in the archaea kingdom and the notes below the charts demonstrate the relative abundance of the most abundant bacteria and the relative abundance of *Geobacter* in the bacteria kingdom.

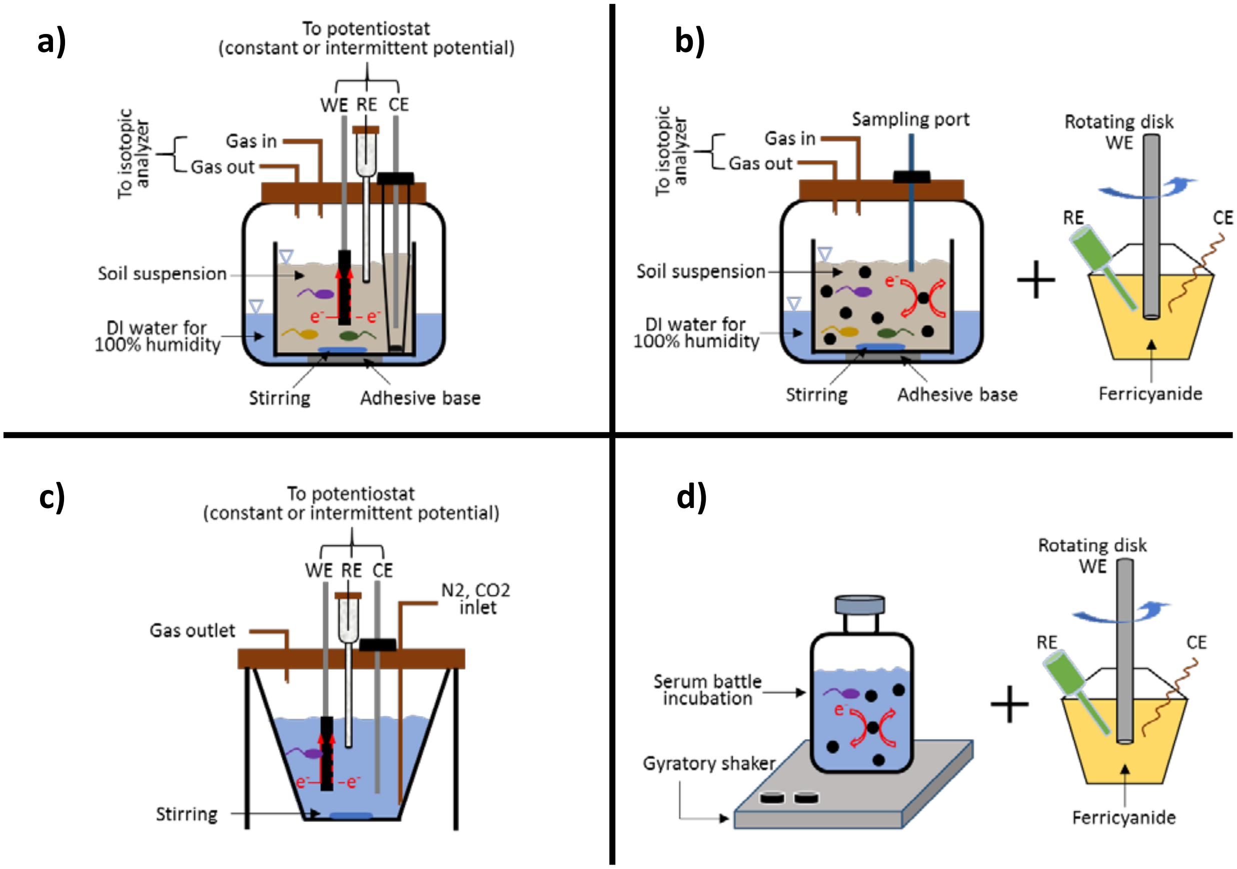


**Figure S3 | Incubation setups. a.** bioelectrochemical peat-soil incubation; **b.** microcosm peat-soil incubation; **c.** bioelectrochemical pure-culture incubation; and **d.** microcosm pure-culture incubation. WE, CE and RE in the figure indicate working electrode, counter electrode and reference electrode, respectively.

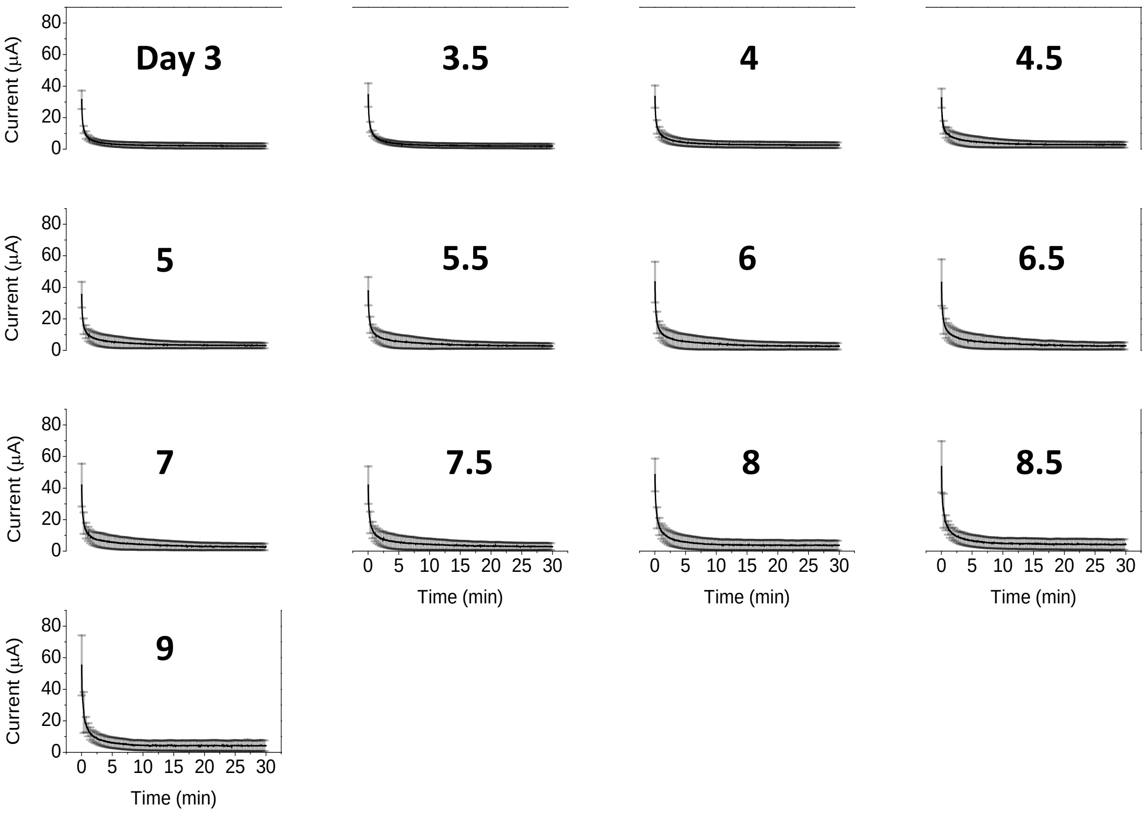


**Figure S4 | Full discharging current profile of the capacitive electron transfer through the pyrogenic carbon matrices during the bioelectrochemical peat-soil incubation.** The capacitive electron transfer started after 3 days of pre-incubation with the conductive electron transfer to facilitate the adaptation of soil microbes to the carbon matrices. The current profile of the first 2 days conductive electron transfer can be found in **Figure 1b** in the main text. The discharging current was recorded every half a day and the discharging duration was 30 min. The recording frequency of the discharging current was 0.1 $s^{-1}$.

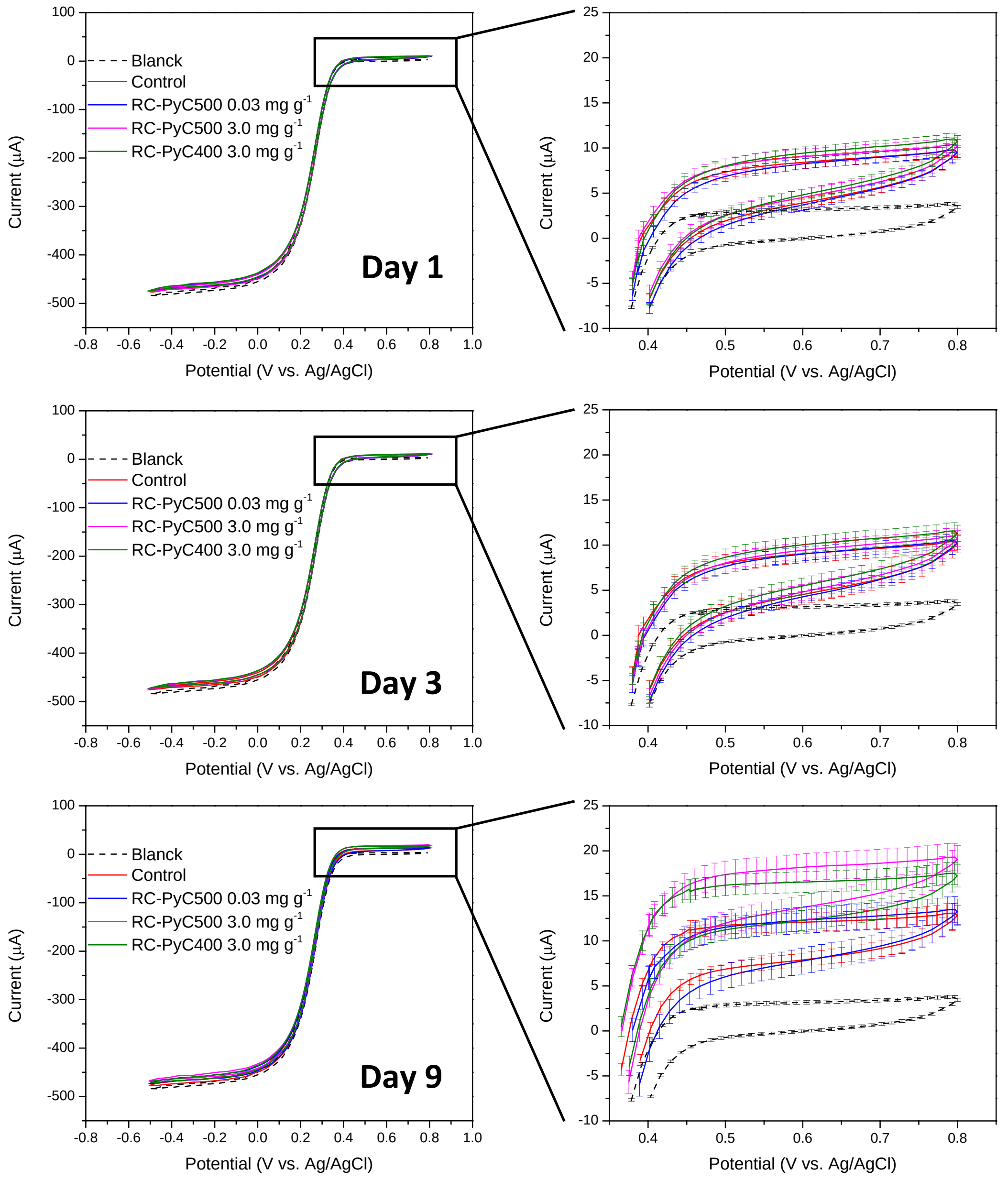


**Figure S5 | Cyclic voltammograms (CVs) of the ferricyanide that was reduced by the peat soil at the microcosm peat-soil incubation day 1, 3 and 9.** Based on the oxidation current increase of ferricyanide (enlarged in the right charts), the number of donated electrons from the peat soil was determined and used to calculate the number of accumulated electrons in the peat soil induced by the redox-cycling electron transfer of the pyrogenic carbon functional groups. In the figure legend, blank indicates the CV of pure ferricyanide without reacting with the peat soil. Control indicates the CV of ferricyanide that was reduced by the peat soil only. RC-PyC500 and RC-PyC400 stand for the CV of ferricyanide that was reduced by the peat soil with the redox-cycling electron transfer (RC) of the pyrogenic carbon (PyC) produced at 500 and 400°C, respectively. The numbers are the application rate of the pyrogenic carbon in the peat soil.

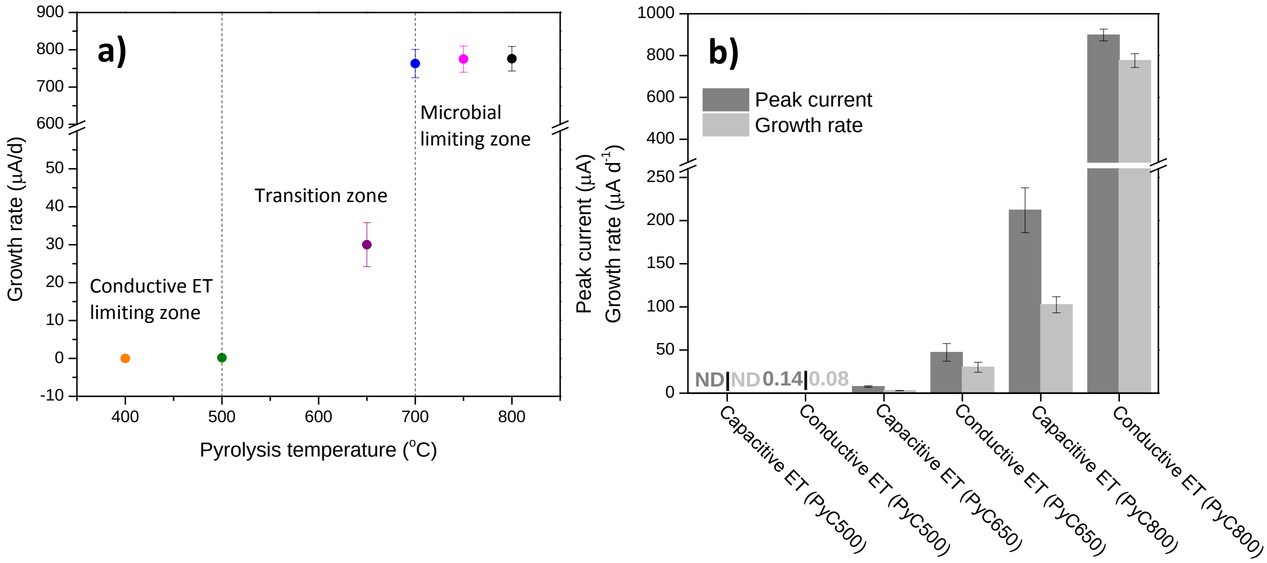


**Figure S6 | Comparison of electron snorkeling kinetics of the pyrogenic carbon matrices in the bioelectrochemical pure-culture incubations. a.** Electron snorkeling kinetics of the conductive electron transfer (ET) in sustaining the alternative respiration of *G. sulfurreducens*. The electron snorkeling kinetics was expressed by the growth rate of *G. sulfurreducens*, which was obtained by the linear fit of the growth current slope (**Figure 3a** in the main text) at the exponential phase. Three kinetic zones had been identified. First was the conductive electron transfer limiting zone (at low pyrolysis temperature range) in which the electron snorkeling was controlled by the conductive electron transfer through carbon matrices; second was the transition zone (at intermediate pyrolysis temperature range); and third was the microbial limiting zone (at high pyrolysis temperature range) in which the electron snorkeling was limited by microbial respiration. That is, even though the conductive electron transfer rate constant still significantly increased about 4 times from $4.8\times10^{-3}$ to $18\times10^{-3}$ cm $s^{-1}$ from the pyrogenic carbon produced at 700°C to that produced at 800°C[24], the electron snorkeling rate remained constant. **b.** Comparison of electron snorkeling kinetics between the conductive and capacitive ET in sustaining the alternative respiration of *G. sulfurreducens*. The comparison was made based on both current peak and current slope at the exponential phase (i.e., growth rate) of the growth current (**Figure 3a** and **b** in the main text) of *G. sulfurreducens*.

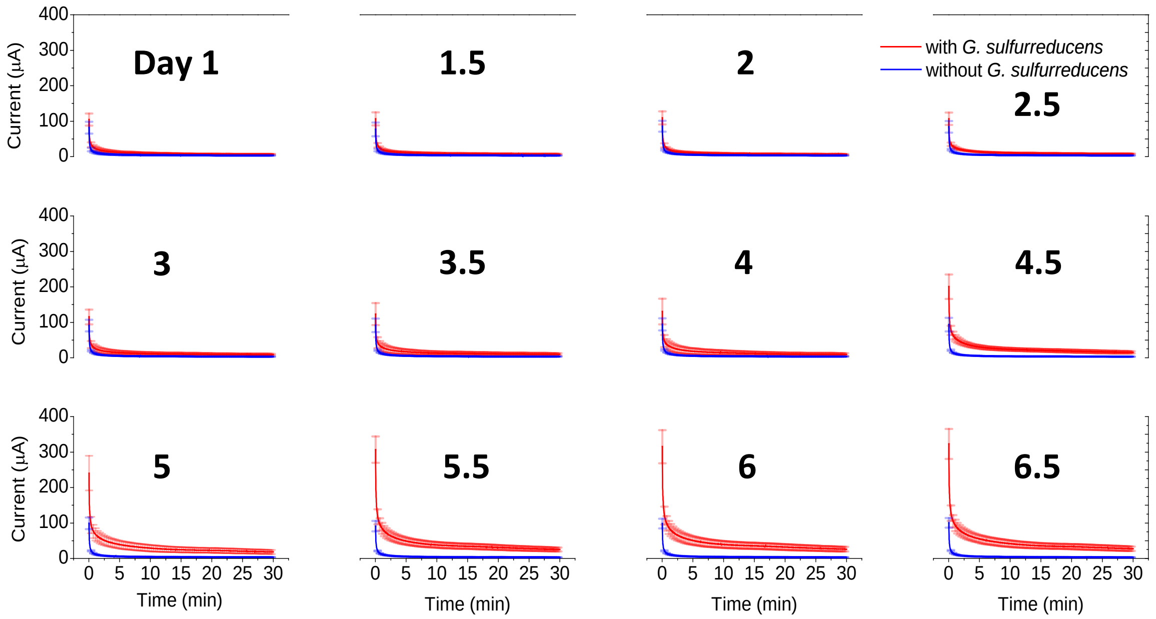


**Figure S7 | Full discharging current profile of the capacitive electron transfer through the pyrogenic carbon matrices that was produced at 800°C during the bioelectrochemical pure-culture incubation.** The capacitive electron transfer started at day 1 of the incubation after 1 day of pre-incubation with the conductive electron transfer to facilitate the adaptation of *G. sulfurreduccens* on the carbon matrices. A control incubation without *G. sulfurreduccens* was also performed to confirm that the increased discharging current through the carbon matrices was a result of the alternative respiration of *G. sulfurreduccens* by using the capacitive electron transfer as an electron snorkeling pathway. The discharging current was recorded every half a day and the discharging duration was 30 min. The recording frequency of the discharging current was 0.1 $s^{-1}$.

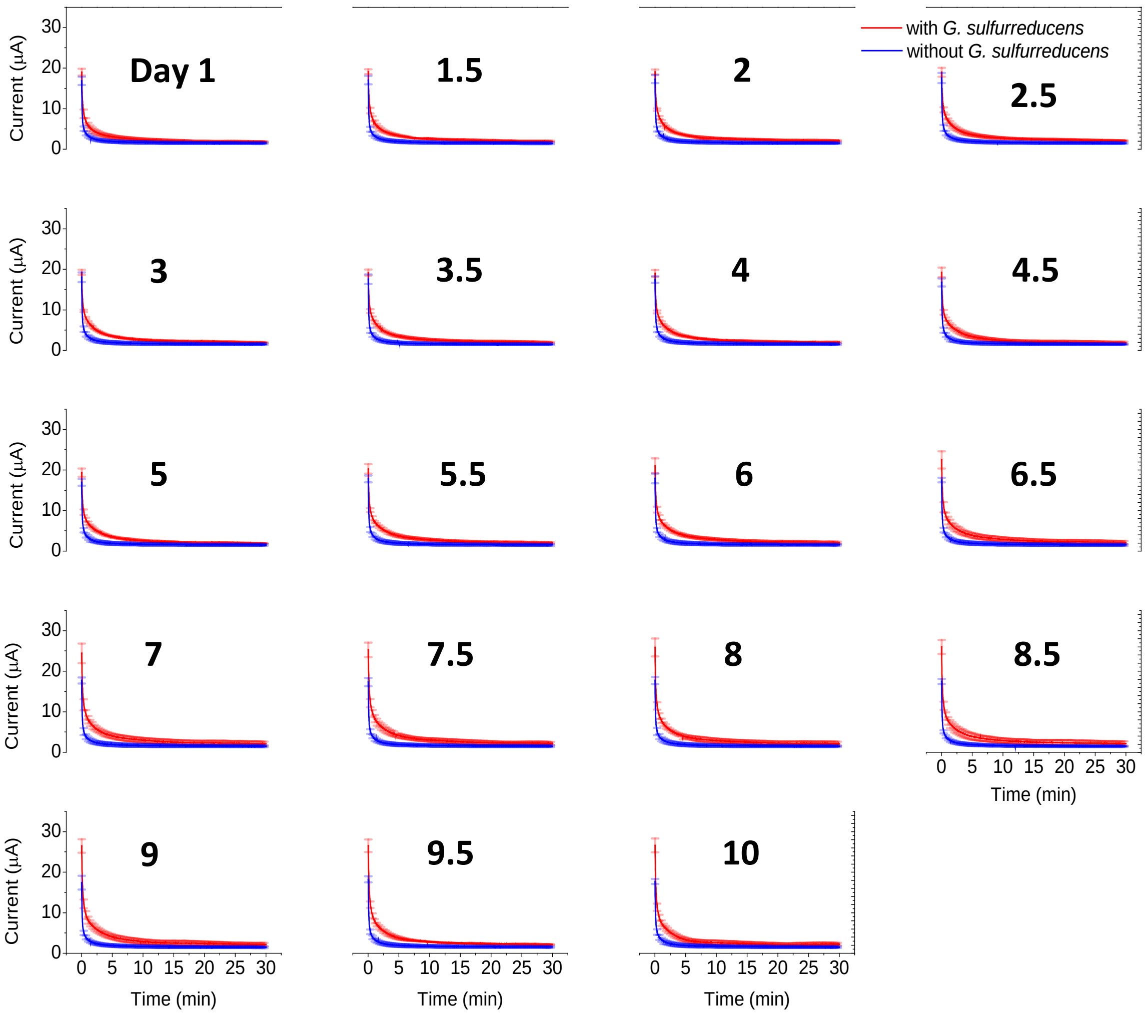


**Figure S8 | Full discharging current profile of the capacitive electron transfer through the pyrogenic carbon matrices that was produced at 650°C during the bioelectrochemical pure-culture incubation.** The capacitive electron transfer started at day 1 of the incubation after 1 day of preincubation with the conductive electron transfer to facilitate the adaptation of *G. sulfurreduccens* on the carbon matrices. A control incubation without *G. sulfurreduccens* was also performed to confirm that the increased discharging current through the carbon matrices was a result of the alternative respiration of *G. sulfurreduccens* by using the capacitive electron transfer as an electron snorkeling pathway. The discharging current was recorded every half a day and the discharging duration was 30 min. The recording frequency of the discharging current was 0.1 $s^{-1}$.

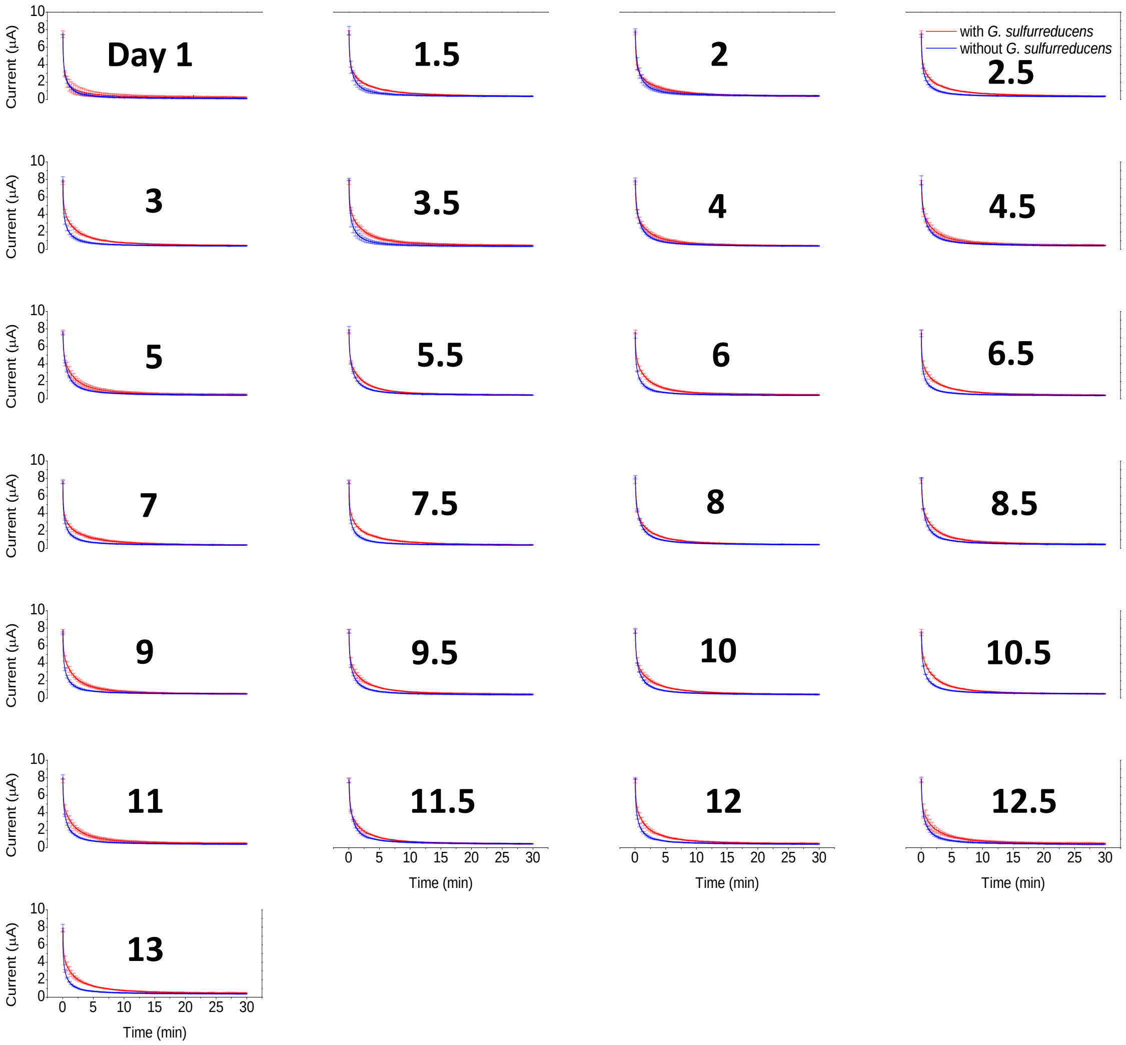


**Figure S9 | Full discharging current profile of the capacitive electron transfer through the pyrogenic carbon matrices that was produced at 500°C during the bioelectrochemical pure-culture incubation.** The capacitive electron transfer started at day 1 of the incubation after 1 day of preincubation with the conductive electron transfer to facilitate the adaptation of *G. sulfurreduccens* on the carbon matrices. A control incubation without *G. sulfurreduccens* was also performed to confirm that the increased discharging current through the carbon matrices was a result of the alternative respiration of *G. sulfurreduccens* by using the capacitive electron transfer as an electron snorkeling pathway. The discharging current was recorded every half a day and the discharging duration was 30 min. The recording frequency of the discharging current was 0.1 $s^{-1}$.

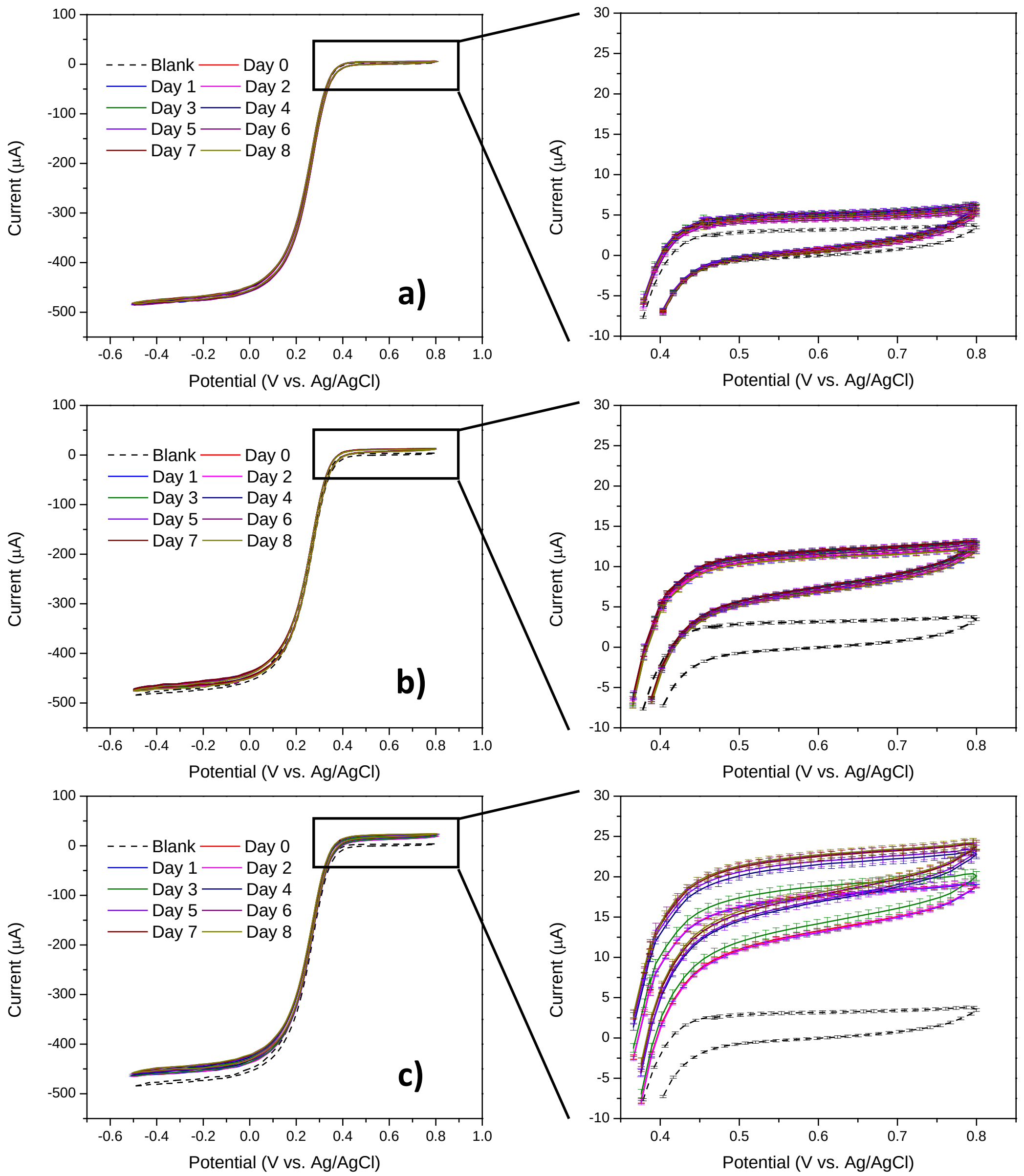


**Figure S10 | Cyclic voltammograms (CVs) of the ferricyanide that was reduced during the microcosm pure-culture incubations.** The oxidation current (enlarged in the right charts) shown in **a** and **b** represents the background reduction of the ferricyanide by only pyrogenic carbon and *G. sulfurreducens*, respectively. In the pyrogenic carbon and *G. sulfurreducens* co-existed incubation (**c**), as microbes growing, surface functional groups started to accept electrons and the pyrogenic carbon became more and more reduced. The reduced pyrogenic carbon was then re-oxidized by donating electrons to oxidizer ferricyanide, which completed the redox cycling and led to an increase of oxidation current in the CVs of ferricyanide. The number of snorkeled electrons shown in **Figure 3c** in the main text was calculated based on the oxidation current in **c** by subtracting the background oxidation current of **a** and **b**.

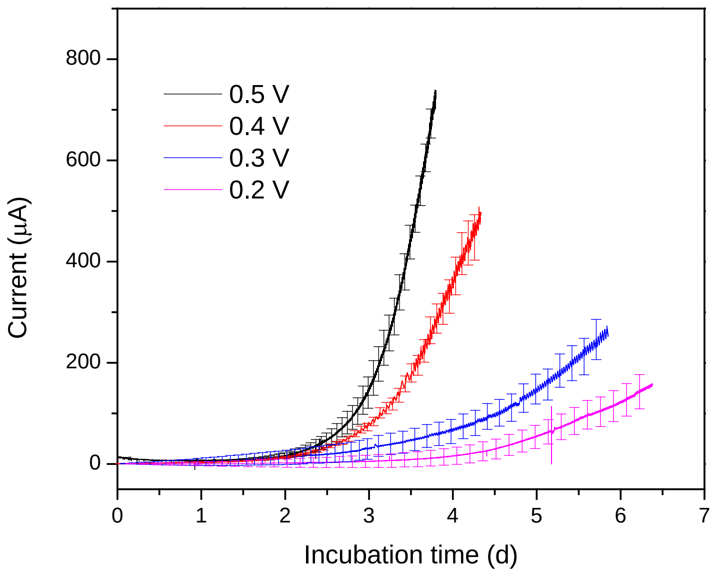


**Figure S11 | Growth current of *G. sulfurreducens* supported by the conductive electron transfer of the carbon matrices during the bioelectrochemical pure-culture incubations.** The growth current was obtained at low to high (+0.2 to +0.5 V vs. SHE) terminal electron accepting potentials to estimate the dependency of the electron snorkeling kinetics on terminal electron acceptors. The electron snorkeling rate was determined by the linear fit of the growth current slope at the exponential growth phase.

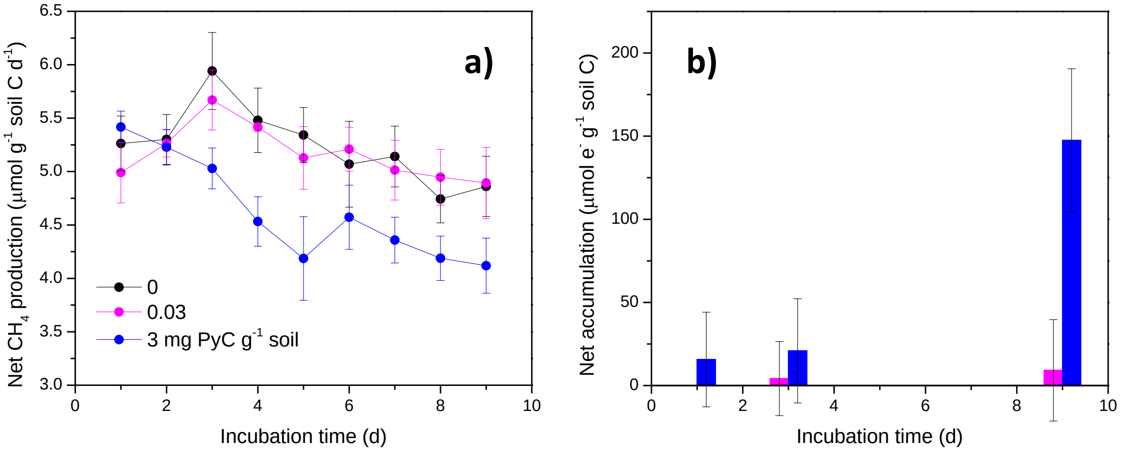


**Figure S12 | Effect of pyrogenic carbon accumulation on $CH_4$ suppression (a) and electron accumulation to support alternative respiration in the peat soil (b).** The chart legend in **a** indicates the concentration of pyrogenic carbon, which increased from 0 to 0.03 to 3 mg pyrogenic carbon (PyC) per gram wet soil. The chart legend in **a** also applies to chart **b.** The net electron accumulation was obtained by setting the electron accumulation in the pyrogenic carbon-free peat soil (i.e., 0 mg PyC $g^{-1}$ soil) as background.

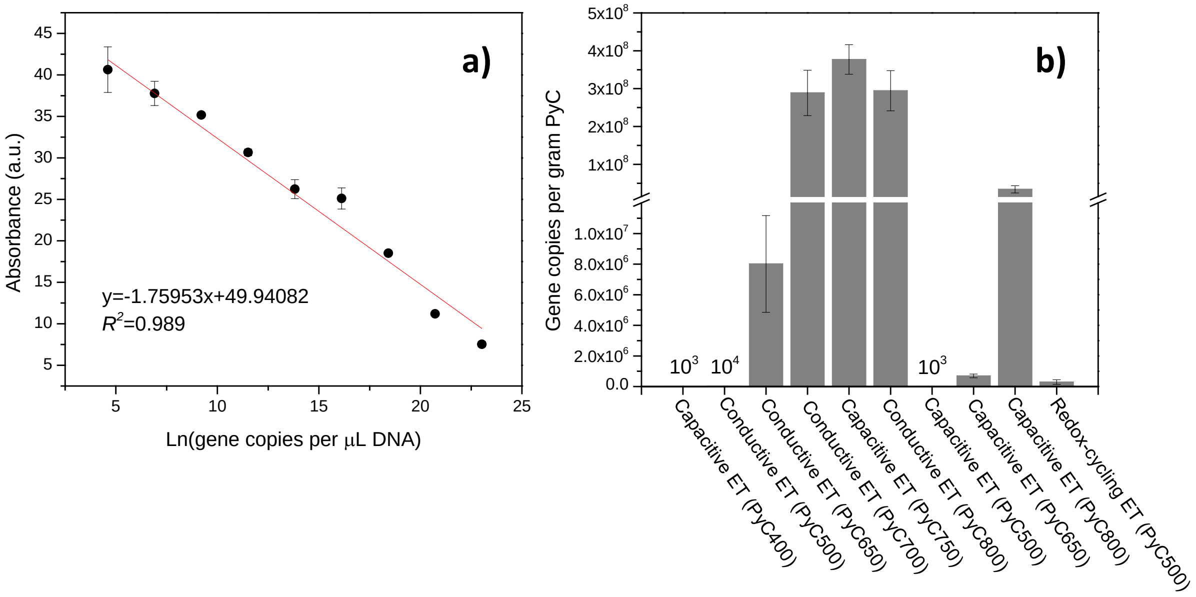


**Figure S13 | Quantification of *G. sulfurreducens* biomass that respires on pyrogenic carbon during bioelectrochemical and microcosm pure-culture incubations. a.** Standard curve. **b.** Copy numbers of 16S rRNA genes of *G. sulfurreducens* respiring on pyrogenic carbon through the conductive and capacitive electron transfer (ET) of the carbon matrices and the redox-cycling ET of the functional groups.

### Supplementary Tables

**Table S1 | $\delta^{13}C$ signatures of the gas phase in bioelectrochemical peat-soil incubations**

| Day | $\delta^{13}CO_2$, Control | $\delta^{13}CO_2$, Conductive ET | $\delta^{13}CO_2$, Capacitive ET | $\delta^{13}CH_4$, Control | $\delta^{13}CH_4$, Conductive ET | $\delta^{13}CH_4$, Capacitive ET |
|---|---|---|---|---|---|---|
| 1 | -34.7±1.2 | -34.7±0.4 | -36.8±1.0 | -44.8±3.6 | -39.1±3.5 | -44.5±2.1 |
| 2 | -37.1±0.6 | -35.4±0.4 | -36.7±0.7 | -40.3±4.1 | -35.3±4.3 | -41.1±4.2 |
| 3 | -37.2±0.4 | -41.1±4.3 | -36.3±0.4 | -36.5±2.2 | -38.2±4.1 | -40.5±6.4 |
| 4 | -38.3±1.8 | -35.1±4.7 | -35.9±0.7 | -42.6±1.1 | -37.4±0.8 | -34.3±4.6 |
| 5 | -37.7±2.2 | -35.6±0.8 | -36.7±0.1 | -45.5±7.8 | -37.5±0.2 | -39.5±4.9 |
| 6 | -35.6±3.5 | -35.2±1.0 | -38.0±0.4 | -38.3±1.3 | -36.8±5.3 | -43.1±2.8 |
| 7 | -35.8±3.1 | -36.7±0.1 | -35.8±2.3 | -39.2±2.6 | -44.4±3.4 | -39.5±3.5 |
| 8 | -35.1±0.4 | -36.6±0.8 | -36.6±0.1 | -38.1±5.8 | -47.5±3.5 | -40.5±4.2 |
| 9 | -36.5±1.6 | -40.2±4.9 | -37.1±2.5 | -41.8±4.2 | -35.5±0.7 | -36.5±4.9 |

**Table S2 | $\delta^{13}C$ signatures of the gas phase in microcosm peat-soil incubations**

| Day | $\delta^{13}CO_2$, Control | $\delta^{13}CO_2$, PyC500-3 mg g-1 | $\delta^{13}CO_2$, PyC400-3 mg g-1 | $\delta^{13}CH_4$, Control | $\delta^{13}CH_4$, PyC500-3 mg g-1 | $\delta^{13}CH_4$, PyC400-3 mg g-1 |
|---|---|---|---|---|---|---|
| 1 | -33.2±1.7 | -20.7±3.8 | -19.5±1.1 | -32.5±3.7 | -28.1±2.8 | -25.1±2.8 |
| 2 | -34.7±0.5 | -22.8±2.5 | -24.7±0.4 | -33.1±2.7 | -27.6±3.4 | -30.4±3.5 |
| 3 | -34.5±0.7 | -23.9±2.9 | -25.8±0.6 | -36.9±2.6 | -33.8±3.6 | -28.6±2.1 |
| 4 | -34.0±0.6 | -24.6±2.8 | -27.3±0.2 | -36.7±2.5 | -28.9±5.5 | -29.2±1.4 |
| 5 | -32.7±0.4 | -24.2±3.1 | -27.1±0.6 | -31.5±2.1 | -29.0±2.8 | -29.5±2.1 |
| 6 | -32.6±2.3 | -26.3±4.2 | -26.6±0.9 | -38.4±6.2 | -33.1±4.4 | -32.3±3.3 |
| 7 | -34.5±0.7 | -27.2±1.3 | -26.9±0.8 | -38.1±5.7 | -33.0±1.4 | -37.5±3.5 |
| 8 | -34.0±0.2 | -28.5±3.5 | -26.7±0.6 | -33.5±3.3 | -34.6±2.0 | -37.4±3.6 |
| 9 | -33.4±0.6 | -29.5±2.1 | -26.5±2.2 | -38.6±2.1 | -35.7±1.8 | -38.7±4.9 |

36 IPCC. *Climate Change 2013: The Physical Science Basis. Contribution of Working Group I to the Fifth Assessment Report of the Intergovernmental Panel on Climate Change*. (Cambridge University Press, 2013).
